# Resolving the 3D landscape of transcription-linked mammalian chromatin folding

**DOI:** 10.1101/638775

**Authors:** Tsung-Han S. Hsieh, Elena Slobodyanyuk, Anders S. Hansen, Claudia Cattoglio, Oliver J. Rando, Robert Tjian, Xavier Darzacq

## Abstract

Chromatin folding below the scale of topologically associating domains (TADs) remains largely unexplored in mammals. Here, we used a high-resolution 3C-based method, Micro-C, to probe links between 3D-genome organization and transcriptional regulation in mouse stem cells. Combinatorial binding of transcription factors, cofactors, and chromatin modifiers spatially segregate TAD regions into “microTADs” with distinct regulatory features. Enhancer-promoter and promoter-promoter interactions extending from the edge of these domains predominantly link co-regulated loci, often independently of CTCF/Cohesin. Acute inhibition of transcription disrupts the gene-related folding features without altering higher-order chromatin structures. Intriguingly, we detect “two-start” zig-zag 30-nanometer chromatin fibers. Our work uncovers the finer-scale genome organization that establishes novel functional links between chromatin folding and gene regulation.

**ONE SENTENCE SUMMARY:** Transcriptional regulatory elements shape 3D genome architecture of microTADs.

## MAIN TEXT

Chromatin packages the eukaryotic genome via a hierarchical series of folding steps ranging from nucleosomes to chromosome territories (*1*). Structural analysis of chromosome folding has been revolutionized by the Chromosome Conformation Capture (3C) family of techniques, which uses proximity ligation of cross-linked genomic loci *in vivo* to estimate contact frequencies (*2*). Interphase chromosome structures such as compartments (*3*), topologically-associating domains (TADs) (*4, 5*), and CTCF/cohesin chromatin loops (*6*) have been characterized using 3C-based methods. Chromosome compartments correspond to large-scale active and inactive chromatin segments and appear as a plaid-like pattern in Hi-C contact maps at the megabase scale (*3*). At the intermediate scale of tens to hundreds of kilobases, topologically associating domains (TADs) spatially organize the mammalian genome into continuous self-interacting domains. TADs are defined as local domains in which genomic loci contact each other more frequently within the domain than with loci outside. TADs appear as square boxes along the diagonal of 3D contact maps (*4, 5*). Mounting evidence suggests that CTCF and cohesin likely mediate TAD formation via a loop extrusion mechanism (*7, 8*) wherein the cohesin ring complex entraps chromatin loci and extrudes chromatin until blocked by CTCF or other proteins. Stabilization of cohesin at CTCF sites result in sharp corner peaks in contact maps and are also referred to as CTCF/cohesin loops or loop domains. Various studies have reported that TADs and CTCF/cohesin loops influence transcription regulation (*9*), and disruption of these structures can lead to certain diseases (*10*). However, the role of RNA polymerase II (Pol II)-mediated transcription on TAD formation and 3D genome organization in general remains controversial, with reports of disparate responses to transcription inhibition (*11–13*).

Currently, CTCF and cohesin are thought to be required for essentially all aspects of genome folding below the level of compartments (*14*). Whether or not the eventual organization has functional consequences remains unclear, as acute disruption of TADs and CTCF/cohesin loops only result in relatively modest effects on gene expression (*15–17*). Beyond CTCF/cohesin-mediated structures, a classical model of transcription posits that *cis*-regulatory elements control gene expression via long-range enhancer-promoter interactions (E-P links) (*18*). Factors such as the general transcription machinery, cofactors, and chromatin remodelers are thought to mediate E-P links that direct transcriptional regulation. Spatial association between the promoters of actively transcribed genes (P-P links) and Polycomb-repressive regions also contribute to “loop-like” conformations (*19*). However, largely due to technical limitations of current techniques, we know little about how these fine-scale structures are organized in the genome, and whether transcription/chromatin factors can mediate the folding of 3D structures. Moreover, gene-specific structures and local nucleosome folding also remain largely unexplored by genome-wide approaches in mammals. The existence of gene loops (*20*), or organization of beads-on-a-string into 30 nm chromatin fibers (*21*), and their functional relevance to transcription regulation, all remain hotly debated. The resolution gap between 1D and 3D genomic maps as well as a paucity of chromatin features below the level of TADs has significantly limited our understanding of gene regulation and its potential link to chromatin architecture. Here, we employed Micro-C, an assay that overcomes the resolution limitations of Hi-C to investigate chromatin organization in mouse embryonic stem cells at a resolution of ~200 bp (i.e. a single nucleosome) (*22, 23*). We focused on dissecting the principles of chromatin folding below the scale of TADs in mammals, in order to probe how transcription and transcription/chromatin regulators may contribute to this more refined scale of 3D genome architecture.

## RESULTS

### Micro-C reveals finer-scale chromatin organizational features in mammalian cells

Micro-C, a method for mapping chromosome folding at length scales from single nucleosome (~100-200bp) to full genomes, has successfully uncovered finer-scale chromatin structures in budding yeast (*22, 23*). To effectively interrogate features below the level of TADs, which are largely inaccessible by Hi-C analysis, we optimized the Micro-C protocol for mapping chromosome folding at single nucleosome resolution in mammalian cells (Fig.1A and fig. S1A-B). We generated 38 biological replicates for mouse embryonic stem cells (mESCs) with sequencing coverages ranging from ~5M to 250M unique reads for each sample. Reproducibility analyses showed high scores (>0.95) across replicates (fig. S1D). We therefore combined all replicates to obtain a total of 2.64B unique reads. In a side-by-side comparison of Micro-C to the current highest-depth Hi-C data (*19*), Micro-C recapitulated all the reported chromatin structures such as compartments, TADs, and loops (fig. S1C), with high reproducibility scores by three independent tests (fig. S1E) and comparable quality scores across various lengths of resolution (fig. S1F). Our results demonstrate that, at coarse resolution, Micro-C yields a qualitatively and quantitatively similar measurement of 3D mammalian genome organization compared to Hi-C analysis.

In addition to the length scale of 100 kb to 100 Mb, corresponding to TADs and compartments (Fig. 1B, blue and gray regions), Micro-C can also assess local chromatin contact behaviors at the scale of 100 bp to 100 kb, as shown by a genome-wide averaged contact frequency analysis (Fig. 1B, pink and purple regions). This range of sizes covers the finer-scale chromatin structures such as genes (median gene length in mouse is ~28 kb), promoter-promoter (P-P) and enhancer-promoter (E-P) interactions (predicted average E-P length is ~24 kb (*24*)), and reveals significantly more loop-like (or “dot”) structures (n=29,548, Material and Methods), which have been largely unresolved by Hi-C analysis (fig. S1G-H). Specifically, Micro-C robustly detects single E-P or P-P linkages with a sharp signal, while standard Hi-C detects weak enrichment, if any, in most of these cases (fig. S1C and H, E-P links).

**Fig. 1.**
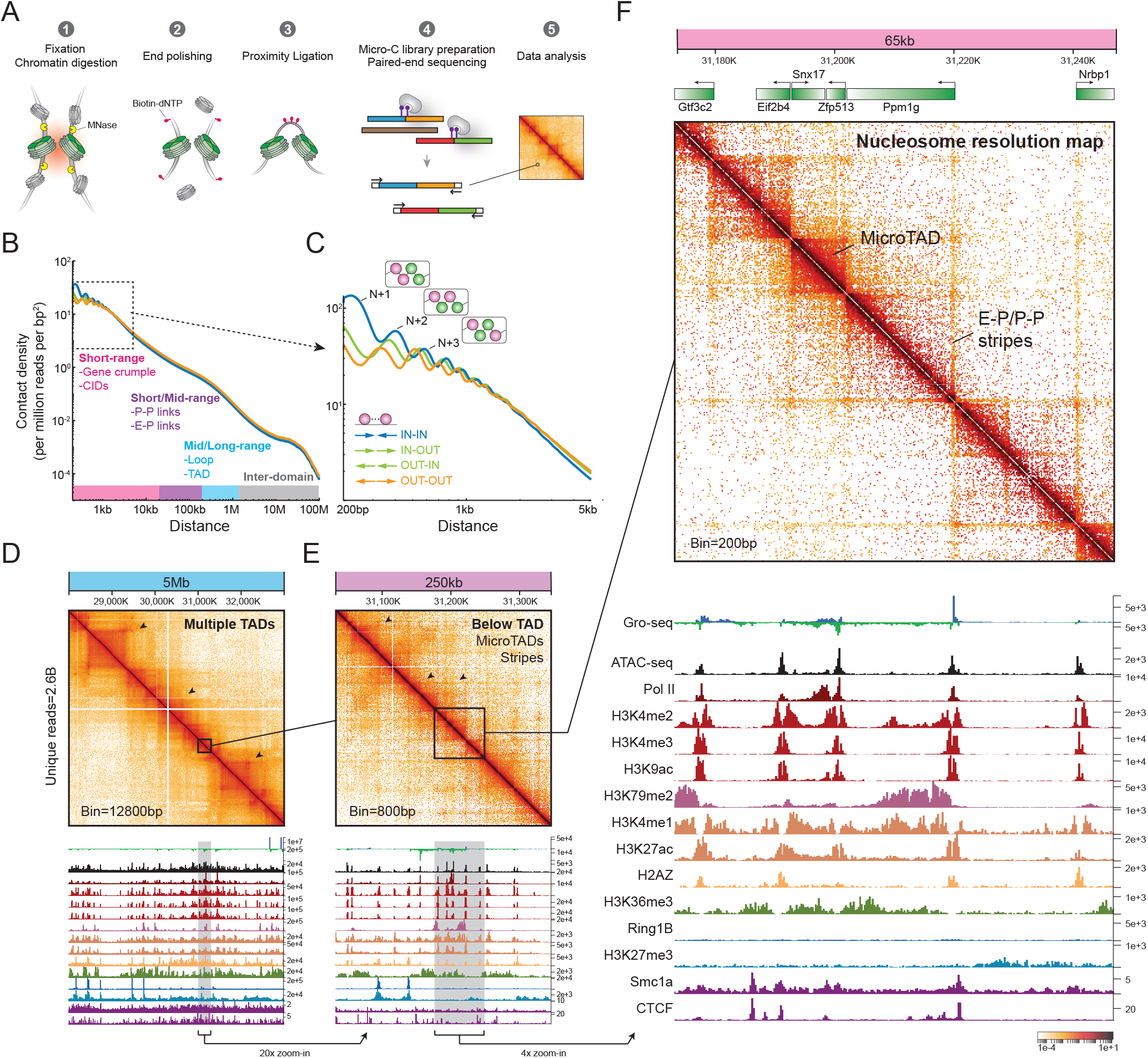
Mapping chromatin folding at single nucleosome resolution. (**A**) Overview of Micro-C method for mammals. (**B**) Decaying curves of inter-nucleosomal contacts with a whole range of genomic distance. Distances along the x-axis are in log10 ratio. Colors coded along the x-axis represent the sizes of chromatin structures at their corresponding scales. Y-axis shows the log10 contact density, normalized to ligated pairs per million reads per bp^2^. The orientations of read pairs facing toward one another are shown separately as IN-IN in blue, IN-OUT and OUT-IN in green, and OUT-OUT in orange (see also fig. S11C). (**C**) Zoom-in of the decaying curve with genomic distance between 200 bp and 5 kb. The first peak represents contact density between adjacent nucleosomes (N/N+1), with data out to ~25 nucleosomes. Schematics illustrate the inter-nucleosomal contacts between N/N+1, N/N+2, and N/N+3, using the tetra-nucleosome for illustration. Ligated nucleosomes are painted in pink. (**D-F**) Multiple zooms of Micro-C contact maps and 1D chromatin tracks on Chr5. Standard white-yellow-red-black heatmap shows the contact intensity for a given pair of bins. (D) A 5-Mb region at 128-kb resolution for TAD visualization. (E) A 250-kb region at 800-bp resolution was 20x zoomed-in from the box in (D). (F) A 65-kb region with gene annotations at single nucleosome resolution was 4x zoomed in from the box in (E). Each dot on the contact matrix represents the contact intensity between a pair of nucleosomes. See fig. S1-S3 for additional examples.

Finally, at the scale of nucleosomal folding (e.g., tetra-nucleosome motifs), Micro-C can capture the contact probability between any given nucleosome (N) and its neighbors (N+1, N+2, N+3, etc.) in all four scenarios of contact directionalities (zoomed plot in Fig. 1C), which allowed us to examine previously undetectable short-range nucleosome folding features such as 30 nm chromatin fibers (discussed below).

### Identification of microTADs and E-P/P-P stripes

Taking advantage of the enhanced Micro-C resolution, we next searched for principles underlying short- and mid-range chromatin folding by integrating our Micro-C contact maps with 48 public genomic datasets (detailed description in Table S1). Beyond TADs, which appears as squares along the diagonal in a 5 Mb region (Fig. 1D), Micro-C also revealed unprecedented details of chromatin organization down to single nucleosome resolution (see 65 kb region in Fig. 1E-F). Visual inspection identified several self-interacting domains spanning ~5-10 kb along the diagonal, encompassing either single genes (e.g., *Eif2b4*), multiple genes (e.g., *Snx17* and *Zfp513*), or intergenic regions (e.g., between the divergent genes *Nrbp1* and *Ppm1g*). Since these domains share various defining features with TADs – they are characterized by higher contact frequency within themselves and reduced contacts outside – but are much smaller than TADs, we will refer to these domains as “microTADs”.

Stripes/flames correspond to lines extending from the diagonal in contact maps and are thought to result from the process of CTCF/Cohesin-mediated loop extrusion (*14*). In Hi-C, stripes are visible at the level of hundreds of kb. But Micro-C uncovered a series of much smaller and nested stripes extending from transcription-start sites (TSSs) and intergenic regions, which colocalize with enrichment of Pol II binding sites, accessible chromatin regions, and active histone marks (Fig. 1F). These stripes do not uniformly correspond to CTCF/Cohesin binding and only extend for ~10-50 kb. Most importantly, these stripes predominantly link promoters and promoters/enhancers together (Fig. 1F and fig. S1C), and are often enriched with sharper signals at their intersections. Thus, these CTCF/cohesin-negative E-P/P-P stripes/links do not appear to simply represent features of the loop-extrusion process, and could be mediated by other proteins and mechanisms.

We confirmed the presence of microTADs and E-P/P-P stripes in mESCs at several other genomic regions containing multiple enhancers that had gone undetected by Hi-C analysis (fig. S2-3). Also, there is no detectable difference in stripe/dot structures between enhancers and “super-enhancers” at these loci (fig. S3). In summary, nucleosome-resolution Micro-C contact maps brings into sharp focus previously obscured chromatin structures within TADs that include microTADs, E-P and P-P stripes, and these finer-scale structures appear to form in specific gene-dependent manner.

### Genome-wide characterization of microTAD boundaries and E-P/P-P stripes

We next analyzed the boundaries of the nested microTADs and E-P/P-P stripes (Fig. 2A). To identify the finer-scale chromatin boundaries in an unbiased way, we used genome-wide insulation score analysis (*25*) and defined boundaries at resolutions ranging from 200 bp to 20 kb. Consistent with previous reports by Hi-C (*26*), TADs can be detected using 10-20-kb resolution boundary metrics (fig. S4A). For example, at 20 kb resolution, we identified 4,384 TAD structures with median size at ~480 kb. Using 200-bp to 1-kb resolution metrics, we identified 361,865 fine-scale boundaries that had previously gone unnoticed (*4*) (Fig. 2A-B and fig. S4A-B) and whose insulation scores precisely peak at the borders of microTADs and E-P/P-P stripes (Fig. 2A). The microTADs have a clear relationship with gene structure, typically encompass one to two genes, and have a median length ranging from 5.4 to 40 kb (Fig. 2C and fig. S4C). In addition, the microTAD boundaries in the active compartments predominantly localize to CpG islands, promoters, and tRNA genes (Fig. 2D), and tend to be closer to the TSS (Fig. 2E), while the boundaries in the gene-poor inactive compartments were found in chromosomal regions rich in repeats (Fig. 2D) and further away from the TSS (fig. S4D). Thus, Micro-C analysis revealed both novel E-P/P-P stripes and allowed high-resolution insulation score metrics establishing the basis for identifying microTAD boundaries.

**Fig. 2.**
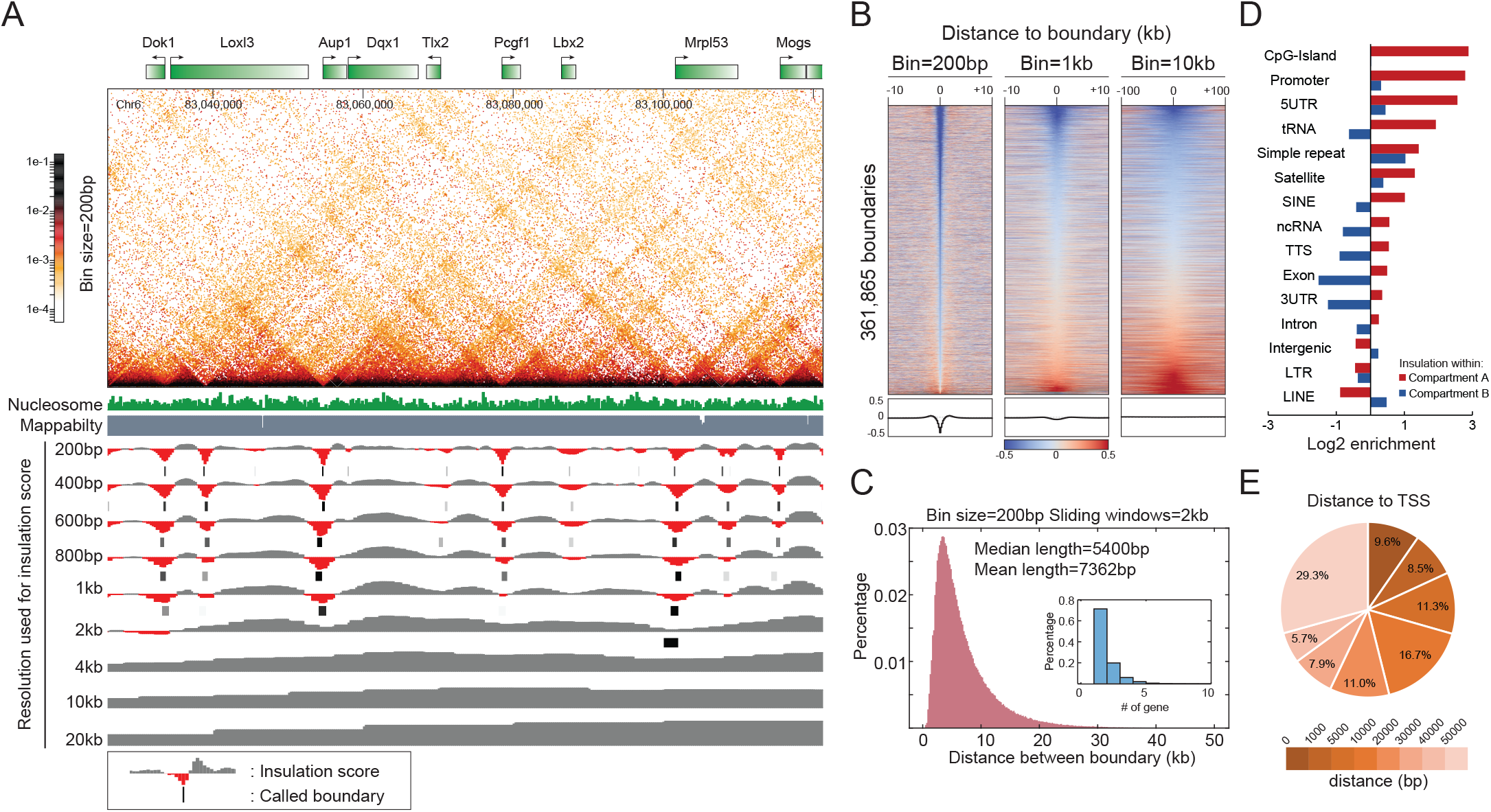
Identification of microTAD boundaries and E-P/P-P stripes. (**A**) Example of microTAD boundary identification. Contact maps were plotted for an 85-kb region on Chr6 at nucleosome resolution. The browser tracks show insulation scores and called boundaries by the data resolution from 200 bp to 20 kb. Called boundaries are indicated as black lines. Nucleosome occupancy and sequencing mappability data shown in the top of browser track indicate that the called boundaries are not artifacts of a low-mappability area or nucleosome-depletion region. (**B**) Heatmap and histogram profile of insulation scores at 200-bp, 1-kb, and 10-kb resolutions. Each row represents insulation scores across ±10-kb region with the called boundary at the center. Diverging blue-gray-red heatmap shows the log2 insulation score for each pixel. Histogram on the bottom of the plot shows the genome-wide average of insulation score for 361,865 boundaries across ±10-kb region of the called boundary at the center. See fig. S4B for additional data. (**C**) Length distribution of microTADs. Distribution of distance between boundaries is plotted in pink, with the number of base pairs in kb for the x-axis, and gene counts in the inset. See fig. S4C for addition data. (**D**) Genomic features of microTAD boundaries. Bar graph shows the log2 enrichment of genomic features at microTAD boundaries at length of 200 bp, with red for boundaries in compartment A and blue for boundaries in compartment B. (**E**) Distance of boundary location to the closest TSS. Pie chart shows the percentage distribution of the distance between the boundary and TSS. Color represents the bin of distance in base pairs.

### Biochemical predictors of microTAD boundary position and strength

The observation that microTAD boundaries tend to overlap with cis-regulatory elements suggests a potential connection between chromatin folding and gene regulation. To further investigate this correlation, we first compared boundary locations with nucleosome occupancy (*27*) (Fig. 3A and fig. S5A-B, Table S1). Overall, the microTAD boundaries are enriched for transcription factor binding sites and dynamic nucleosomes (also known as “fragile nucleosomes”) (Fig. 3A and fig. S5A-B), since signals from short fragments (1-100 bp) corresponding to transcription factor binding are substantially higher at these boundaries. Notably, boundary strength is also correlated with nucleosome occupancy (fig. S5C), as stronger boundaries exhibit a higher level of nucleosome depletion and often center at nucleosome-depleted regions (NDRs).

**Fig. 3.**
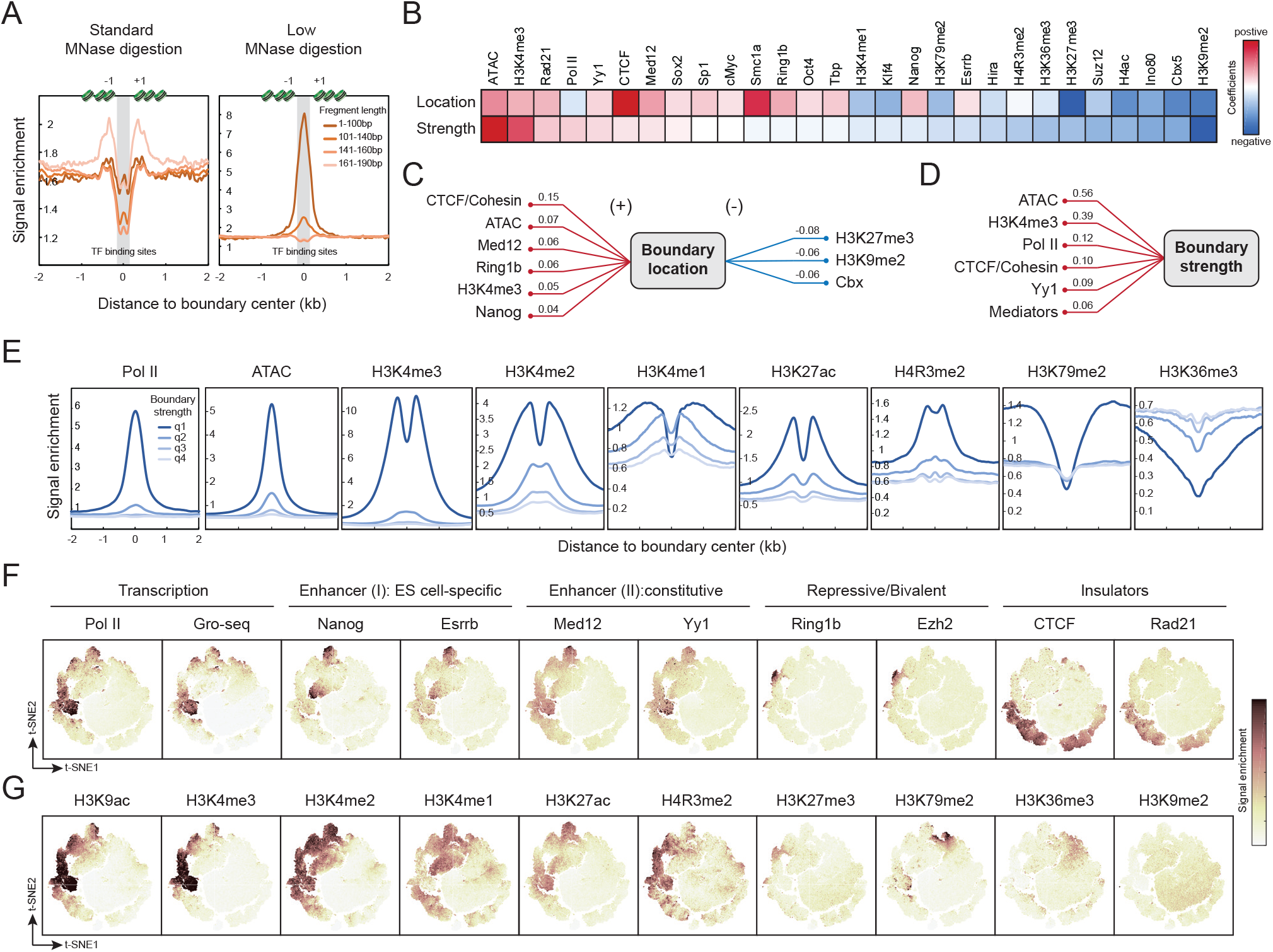
Features of microTAD boundaries. (**A**) MicroTAD boundaries enrich for the dynamic/fragile nucleosomes and transcription-binding sites. Nucleosome occupancy measured by MNase-seq mapping was plotted as signal enrichment for y-axis against x-axis with ±2-kb distance from the microTAD boundary. Fragment length at 161-190 bp in the standard MNase digestion represents nucleosome occupancy, and 1-100-bp fragments in the low-level MNase digestion represents transcription factor binding sites. (**B**) Predicting boundary location and strength by genome-wide data. Heatmap shows the generalized linear regression coefficient of factors for microTAD boundary predictions. (**C-D**) Key parameters to predict boundary location and strength are highlighted as a wiring chart in numeric form. (**E**) MicroTAD boundaries enrich for active marks. Histogram shows the signal enrichment on the x-axis and ±2-kb region of the microTAD boundary on the y-axis. Signal enrichments across the regions were sorted according to quartiles based on the boundary strength. See additional examples in fig. S5G-H (**F-G**) MicroTAD boundaries consist of a heterogeneous population. Boundaries were plotted in a 2D space by the t-SNE score. Heatmap was coded by the signal intensity of the target in question. See additional data in fig. S6B.

Given that the majority of microTAD boundaries are flanked by +1 and −1 nucleosomes immediately upstream and downstream of a gene’s TSS (Fig. 3A and fig. S5A), we next asked how local histone modifications and protein binding profiles might relate to microTAD boundary properties. To systemically identify factors most predictive of microTAD boundary location and/or strength, we applied a generalized linear regression model based on genome-wide chromatin data (fig. S5E). Key predictors (28 out of 48, Table S1) were obtained after removing redundant factors using lasso regularization. The regression coefficients revealed that transcription factors, architectural proteins, and repressive regulators are the most predictive factors of microTAD boundary properties to varying degrees (Fig. 3B). For example, CTCF/Cohesin, chromatin accessibility, Mediator, active histone marks and transcription factors are all positive predictors of boundary location, while H3K27me3 and H3K9me2, etc. are also predictive, but with negative regression weights (Fig. 3C and fig. S5F). Strikingly, CTCF/Cohesin are the best predictors of boundary location but only moderate predictors of boundary strength. Instead, chromatin accessibility and transcriptionally active chromatin predict boundary strength better than CTCF/Cohesin occupancy (Fig. 3D).

To further explore the relationship between chromatin features and microTAD boundary strength in a more quantitative manner, we sorted boundaries into four quartiles (q1 to q4) with decreasing insulation strength and calculated signal intensities around them for several candidates. This confirmed that microTAD boundary strength positively correlates with the signal intensity of the predictive factors identified above (Fig. 3E and fig. S5G-H). Histone modifications on the boundary-spanning nucleosomes also correlate with the level of boundary strength (Fig. 3E and fig. S5G). Overall, active marks such as H3K4me1/2/3, H3K9ac, and H3K27ac are all highly enriched on nucleosomes flanking a strong boundary, while repressive (H3K27me3, H3K9me2) and elongation (H3K36me3) marks anti-correlated with the boundary activity. Thus, unlike TADs which appear to be demarcated mainly by CTCF/cohesin (*6*), microTADs show a wide spectrum of boundary markers.

### A combination of chromatin features discriminate subgroups of microTAD boundaries

We next sought to examine whether novel combinatorial chromatin patterns can segregate microTAD boundaries into subgroups. To dissect the properties of a boundary, we plotted single boundaries in 2D t-Distributed Stochastic Neighbor Embedding (t-SNE) space according to their protein-binding profile (fig. S6A-B). We can classify the microTAD boundaries into at least five partially overlapping subgroups based on their biochemical and functional features (Fig. 3F). We denote the five boundary subgroups: 1) transcription-dependent; 2) enhancer-related (ES cell specific); 3) enhancer-related (constitutive); 4) repressive; and 5) CTCF/Cohesin-mediated.

Transcription-dependent boundaries are strongly enriched for active features such as Pol II, suggesting that transcription may be a key driver of fine-scale chromatin folding. A subset of H3K4me3-enriched boundaries also showed elevated H3K27me3 signal, a bivalent chromatin mark of certain developmental genes in stem cells (*28*). Interestingly, enhancer-related boundaries occupy two distinct t-SNE spaces, one enriched with H3K4me1, Esrrb, and Nanog (Enhancer I), and another enriched with H3K27ac, mediator, and Yy1 (Enhancer II). Our results suggest that Enhancer I boundaries might be cell-type specific, while the Enhancer II group includes constitutive boundaries across cell types. In general, active promoters and cis-regulatory elements can form boundaries that spatially segregate kb-sized regions into distinct microTAD domains, and at least one subgroup of the domains appears to be cell type-specific.

Strikingly, CTCF/Cohesin binding describes a subgroup of boundaries clearly separated from the other subgroups in t-SNE space, implying that CTCF/Cohesin boundaries mostly contribute to chromatin architecture, while their spatial associations appear to be separated from the transcription and enhancer classes. Finally, histone modifications not only categorize the majority of the microTAD boundaries (Fig. 3G), but also identify some rare populations. For instance, H3K79me2-enriched boundaries might be related to DNA replication and repair (*29*). We conclude that microTAD boundaries are heterogeneous and fall into distinct subclasses, suggesting that multiple distinct mechanisms can contribute to microTAD formation.

### Transcription factors and CTCF/cohesin dictate chromatin architecture within TADs

We next analyzed how microTADs, E-P/P-P stripes, and CTCF/cohesin loops interact with each other to form nested structures at scales of tens to hundreds of kilobases (fig S7A-B). The stripes extending from microTAD boundaries connect promoters to promoters (P-P links) (Fig. 4A and fig S7A-B, red arrows), enhancers to promoters (E-P links) (Fig. 4B and fig S7A-B, green arrows), CTCF/Cohesin binding sites (CTCF/Cohesin loops) (fig S7A-B, purple arrows), and even TSS to TTS within the same gene (gene loops) (Fig. 4A and fig S7A-B, pink arrows). Interestingly, previous studies reported that Polycomb repressive regions interact with each other through “loop” structures (*19, 30*). However, instead of forming focal contact enrichments, we found that H3K27me3-rich regions form nested sets of chromatin contacts in Micro-C maps, as one patch of repressive chromatin hooks-and-loops to another patch (Fig. 4C and fig. S7C, blue arrows). These bundle interactions are often delimited by a broad region of H3K9me2 (Fig. 4C and fig. S7C, gray arrow). Thus, whereas polycomb interactions appear as blobs or loops in Hi-C, the higher resolution of Micro-C resolves the ultrastructure of these interactions and reveals them to occur as nested sets.

**Fig. 4.**
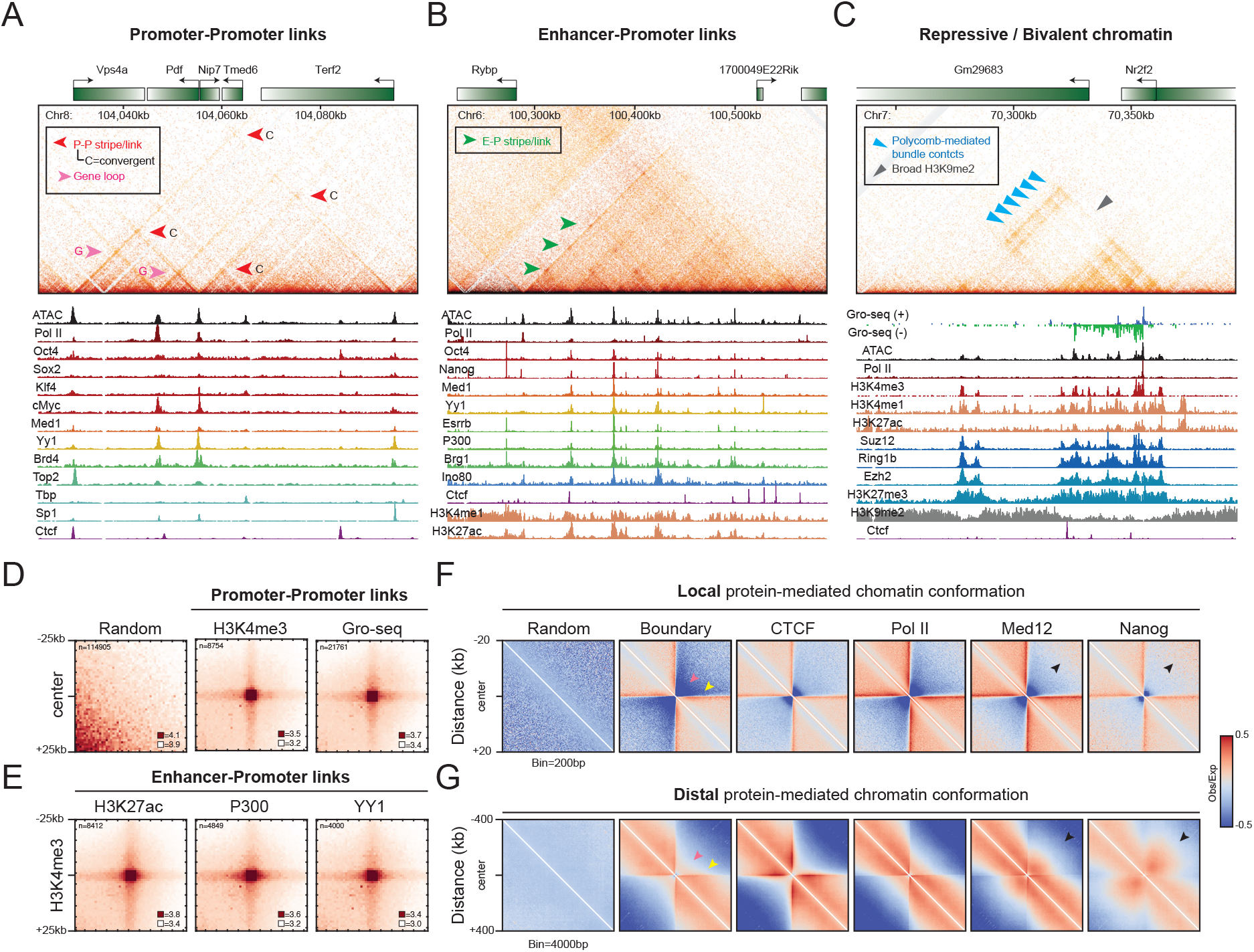
Protein-centered fine-scale chromatin folding. (**A-C**) Examples of promoter-promoter links, enhancer-promoter links, and repressive chromatin patches, also see fig. S7 for additional examples. Contact maps were shown in 100-bp resolution. Red arrow: P-P stripe/link. Pink arrow: Gene loop, or TSS-TES stripe/link within a gene. Green arrow: E-P stripe/link. Blue arrow: Polycomb-repressive bundle contacts. Gray arrow: Broad H3K9me2-rich region. Since we cannot exclude that the promoter/gene body of one gene serves as an enhancer for another gene, the terms P-P and E-P links are necessary ambiguous in some cases unless there is mechanistic evidence in support of one or the other at any given locus. (**D-E**) Protein-mediated loop/dot structure. Pile-up analysis of dot enrichment was plotted according to the loci of paired ChIP-seq peaks, including the canonical marks for P-P and E-P interactions. Loci with CTCF peaks bound within ±5 kb were removed to reduce signals contributed by CTCF loops. See additional data in fig. S8A-B. (**F-G**) Pile-up analysis of the target-centered chromatin structure. Maps were plotted by 200-bp resolution data for local chromatin folding or by 4000-bp resolution data for long-range chromatin organization. Contact matrix was normalized to distance, with positive enrichment in red and negative signal in blue. Pink arrow marks microTAD boundary and yellow arrow marks protein-associated stripes. Black arrows indicate the strong interaction off the diagonal on the maps. See additional examples in fig. S8C.

The direction of Pol II elongation (or gene orientation) is one of the major determinants of chromatin folding in yeast (*23*). We thus asked whether gene orientation also controls fine-scale chromatin folding in mammals. Overall, we found that P-P stripes usually extend toward the same direction as Pol II elongation, and link the promoters of co-regulated genes (Fig. 4A-B and fig. S7A-B). We observed three major types of P-P stripes: convergent, tandem, and divergent stripes. Convergent stripes emanating from a pair of convergent genes collide with each other and form a strong signal at the cross point (Fig. 4A and fig.S7A, “C arrows”). When a stripe extends through a series of tandem genes, we observe higher contact frequencies between their TSSs, suggesting that DNA links sometimes connect tandem-gene promoters (fig.S7A, “T arrows”). While the majority of divergent promoters form a strong boundary (fig. S7D), pairs of adjacent divergent genes are sometimes linked together by P-P stripes, suggesting that antisense transcription or arrested Pol II might also mediate P-P links (fig.S7A, “D arrows”). E-P links are structurally similar to P-P links, with a stripe extending from a promoter and connecting to its distal enhancer element(s), marked by high levels of H3K4me1, H3K27ac, mediator, P300 and pluripotency transcription factors (Fig. 4B and fig. S7A-B, green arrows). Genome-wide analysis of paired loci further revealed ubiquitous dots linking some promoters to other promoters (Fig. 4D), enhancers to promoters (Fig. 4E), or genomic loci bound by transcription/chromatin factors (fig. S8A-B). These dots are usually independent of CTCF/cohesin binding sites, suggesting that cis-regulatory elements and their associated proteins may also significantly contribute to loop/dot formation with varying degrees of strength.

We next explored how different proteins regulate local and distant *cis*-interactions at the genomic scale. Low (4-kb) and high (200-bp) resolution 2D pile-up contact maps not only revealed distinct chromatin structures at promoter/enhancer factor-binding sites but also confirmed that these loci generally colocalize to the regions enriched with boundaries and stripes (Fig. 4D-E, pink and yellow arrows). Specifically, CTCF/cohesin binding sites colocalize with strong distal boundaries, and are associated with loop-extrusion stripes extending up to ~300 kb (Fig. 4D-E and fig. S8C) (*31*). In contrast, Pol II and active histone modifications mark robust short-range boundaries but are associated with weaker long-range insulators compared to CTCF/cohesin (Fig. 4D-E and fig. S8C). Evident short-range stripes extend beyond Pol II-occupied sites, likely a result of combining gene and P-P/E-P stripes. Interestingly, contact maps centered at enhancers (i.e., sites occupied by mediator, pluripotency factors like Nanog and cofactors like P300) show distinctive interactions connecting their upstream and downstream genomic loci at lengths between 10 and 100 kb (Fig. 4D-E and fig. S8C, black arrows). We speculate that a single enhancer may contact one to multiple genomic loci and keep them in spatial proximity, which can be detected by Micro-C. Together, this analysis suggests that transcription-associated factors form stronger insulating boundaries at short range, but that CTCF and cohesin form boundaries that insulate more strongly at longer range.

### Pol II-dependent transcription drives gene folding

We previously reported that gene-specific compaction levels are anti-correlated with transcription rate in the budding yeast *Saccharomyces cerevisiae* (*22*). This suggested that active transcription influences the unfolding of genes, and vice versa (*32*). Does this also hold in mammals? We first plotted the density of Micro-C counts within each gene against its Pol II enrichment (Fig. 5A). Surprisingly, in stark contrast to the findings in budding yeast, gene compaction levels are positively correlated with transcription activity (Spearman’s Rho=0.6). We corroborated this result by comparing gene folding levels by both unscaled (Fig. 5B) and scaled (fig. S9A) pile-up analyses. Interactions are notably more pronounced within highly-transcribed genes and genes with elevated levels of bound Pol II (Fig. 5B and fig. S9A-C) compared to lower-expressed genes. In addition to gene folding, P-P links also correlate with transcription activity (Fig. 5C and fig. S9D). The TSSs of highly transcribed genes often contact others with high transcription rates, while the TSSs of weakly transcribed genes show a decreased contact probability to other TSSs. Several P-P links are formed even in the absence of a CTCF/cohesin binding site nearby (± 5 kb, Fig. 5C). These results reveal that the levels of gene folding and P-P links correlate with the transcription activity at these loci.

**Fig. 5.**
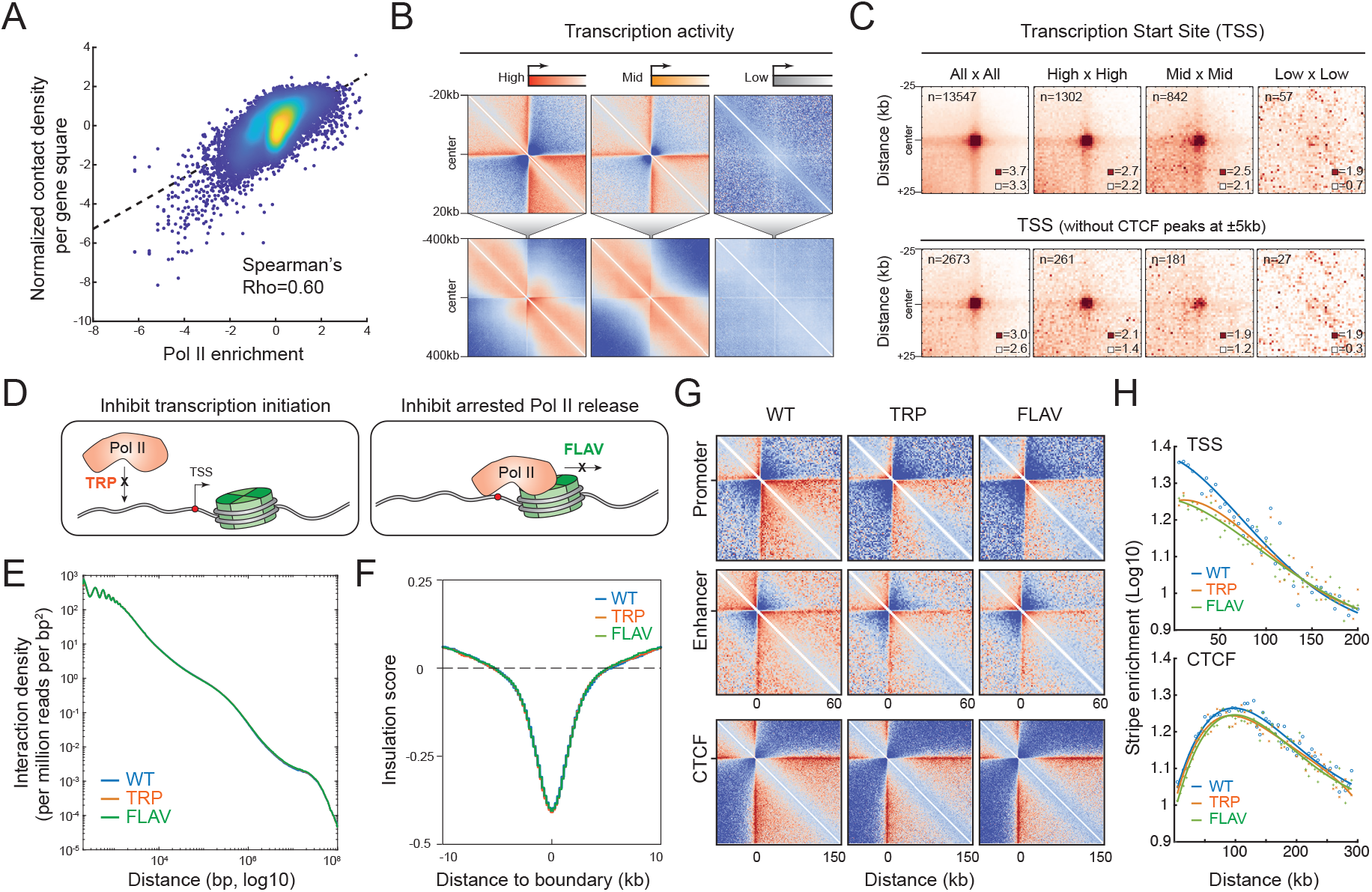
Transcription-dependent gene folding. (**A**) Scatter plot of gene compaction with transcription activity. X-axis shows Pol II ChIP signal enrichment, and y-axis shows contact density per gene square surface, bp^2^. Contacts per gene were quantified by Micro-C read pairs over 300 bp. (**B**) Pile-up analysis of TSS-centered chromatin structure for local (top) and distal (bottom) structures. The contacts in the actively transcribed genes are the sum of the promoter interaction with the gene body (manifested as a stripe extending from the TSS) and intra-gene contacts. Contact maps are shown in log10 contact counts normalized by matrix balancing or by distance. (**C**) TSS-mediated chromatin links. Genome-wide averaged contact maps were plotted according to the loci of paired TSSs, with CTCF or without CTCF within ±5 kb of the TSS. (**D**) Schematics describing the mechanism of Pol II inhibition. In this study, flavopiridol and triptolide treatments were performed at 1 µM final concentration for 45 min. (**E**) Whole-genome contact density decaying curves. Pol II inhibitions have little effect on global chromatin folding, as the three curves overlap entirely. (**F**) Histogram of genome-wide averaged insulation score. Pol II inhibition has no global effect on microTAD boundary strength. (**G**) Pol II inhibitions block gene and enhancer stripes but not CTCF stripes. Target-centered pile-up maps were plotted by the 1-kb resolution data and normalized to distance. (**H**) Signal decaying curve of the gene stripe. X-axis shows the normalized contact of gene stripe, and y-axis shows the distance to the target center up to 300 kb.

To unravel the functional relationship between gene structure and transcriptional activity we acutely inhibited transcription by treating cells with either triptolide or flavopiridol, drugs that inhibit promoter melting and productive elongation, respectively (Fig. 5D). Acute inhibition of transcription had little effect on global chromatin organization at the scale of compartments, TADs, and loops (Fig. 5E and fig. S10A). Pol II inhibition also did not affect the strength of microTAD boundaries (Fig. 5F and fig. S10B-C). This result is consistent with the previous observation that DNA binding transcription factors and CTCF/cohesin may be sufficient to maintain the global chromatin configuration in mammalian cells, as Pol II inhibition also had a negligible effect on chromatin organization in the fly embryo (*11*). However, the intensities of gene stripes significantly decreased upon Pol II inhibition, and enhancer stripes became moderately reduced, while CTCF/cohesin stripes remained largely unaltered (Fig. 5G-H). These findings suggest that Pol II drives chromatin folding at the gene level but has little or no effect on higher order chromatin organization.

### Secondary chromatin folding in mammals

We next explored potential second order chromatin folding, popularly known as the 30 nm fiber. Dominant models for 30 nm fibers include the one-start solenoid helix model and the two-start zig-zag model, which differ in their periodicity, nucleosome spacing, and linker histone binding (fig. S11A) (*21*, *33*–*36*). However, evidence for existence of a 30 nm chromatin fiber *in vivo* has been elusive. A few studies have characterized local nucleosome folding by imaging or genomic approaches, finding some evidence of a tri- or tetra-nucleosome motif in yeast (*22*) and two-start helical fibers in human (*37, 38*).

We thus interrogated Micro-C data to probe for evidence of 30 nm chromatin fibers in mammalian cells (Fig. 6A-C and fig. S11B-F). Strikingly, we observed a unique pattern of nucleosomal folding in mouse cells that supports the two-start zig-zag model (or zig-zag tetra-nucleosome folding) (*39, 40*). The abundance of ligation products for N/N+2 is nearly identical to N/N+3, and N/N+4 is also similar to N/N+5, and so forth up to ten nucleosomes away (Fig. 6B-C and fig. S11E-F, black arrows), consistent with prior Cryo-EM results (*41*). In contrast, the slopes in the yeast data appear to monotonically decrease without apparent periodic patterns (Fig. 6B-C and fig. S11E-F). Therefore, we conclude that an extended zig-zag path of chromatin folding can be detected in mES cells, which may consist of at least 2 – 3 tetra-nucleosome stacks (Fig. 6C), while yeast secondary chromatin folds into a loose zig-zag-like structure that contains sparse tri-or tetra-nucleosome motifs.

**Fig. 6.**
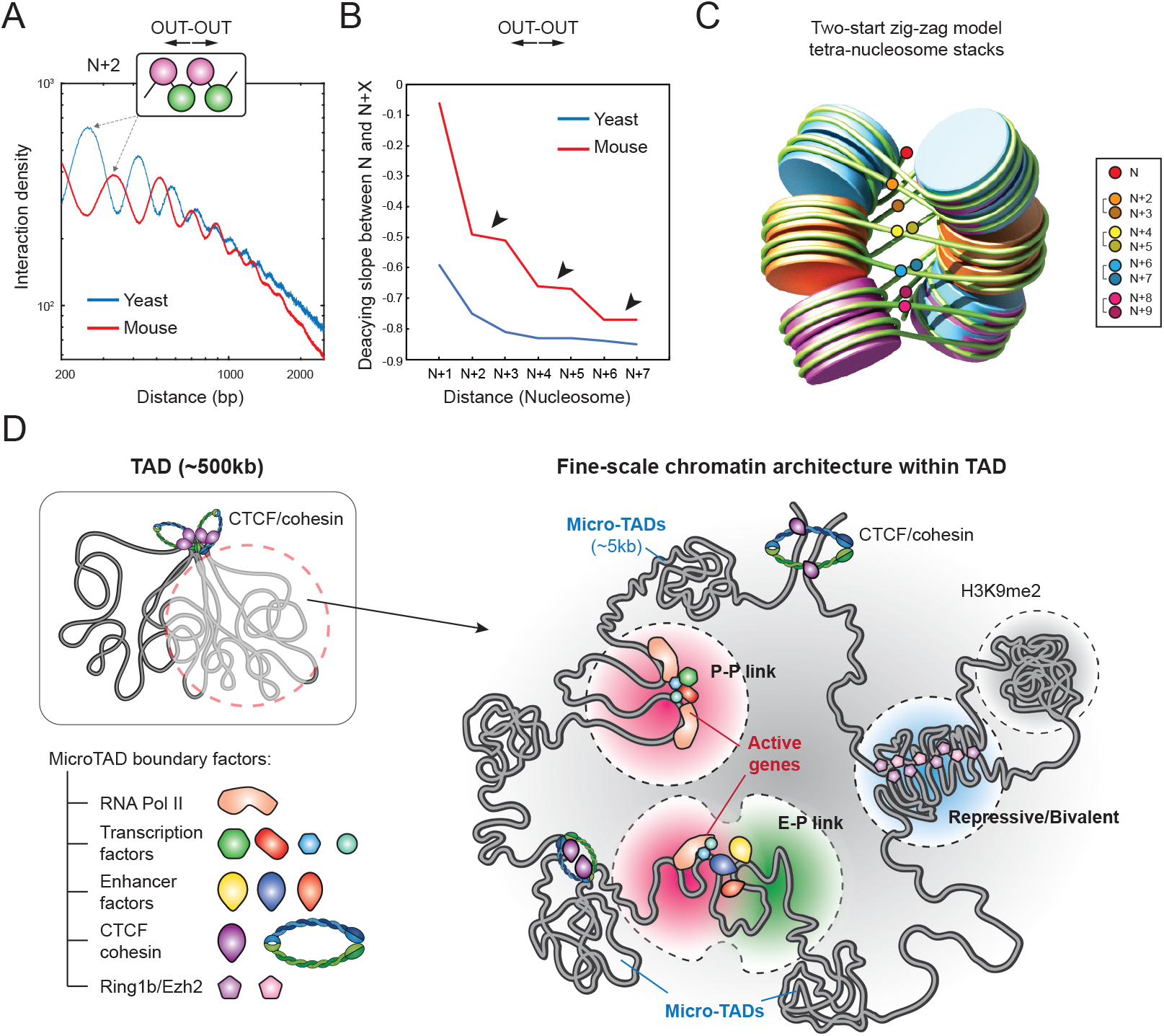
Models of the fine-scale chromatin folding. (**A**) Interaction decaying curves of mouse and yeast Micro-C data. Here we only showed the ligation products in “OUT-OUT” orientation. Distances along the x-axis are from 200 to 2000 bp in log10 ratio, and y-axis shows the log10 contact density, normalized to ligated pairs per million reads per bp^2^. (**B**) Interaction abundance of N/N+X with distance. Curve indicates the slopes between the peak point of N+2 and N+X in Fig. 6A. X-axis represents the distances in units of nucleosomes, and y-axis is the slope of the indicated N/N+X. Black arrows highlight that every two adjacent nucleosomes (e.g. N+2 and N+3) exhibit a similar level of interactions to nucleosome N. (**C**) The two-start zig-zag tetra-nucleosome stacks. The colored dots represent the ligated partners between N/N+X in the “OUT-OUT” orientation. (**D**) Schematics of the new layer of chromatin organization revealed by Micro-C. The fine-scale chromatin structures such microTADs, E-P/P-P links, and repressive bundle contacts build the basic units within TADs.

## DISCUSSION

How much the spatial organization of mammalian genomes into compartments, TADs, and loops actually impacts gene regulation remains hotly debated (*42, 43*). Recent findings suggest that genome topology and transcription are only loosely linked at the level of TADs and loops (*11*–*13*, *15*–*17*, *19*), suggesting that there may be an additional layer of structures that more directly relate to gene regulation. One of the key missing pieces to connect 3D genome architecture to transcription has been the inability of Hi-C to attain the resolution necessary to discern gene-level features of chromatin. Here, we describe a method, Micro-C that dissects chromosome folding in mammals at scales from single nucleosomes to the whole genome. Our high-resolution Micro-C contact maps have revealed several novel features that justifies an updated model for 3D genome organization at the resolution relevant to gene regulation (Fig. 6D). Micro-C reveals that E-P/P-P links act as a hub to connect gene and cis-regulatory elements while also demarcating regions below the level of TADs into distinct chromatin domains we call microTADs.

Micro-C now shows that E/P-P/P interactions often manifest as stripes extending from the borders of microTADs and that these stripes can link multiple genes or genes and enhancers together. We propose three mechanisms that might mediate the formation of E-P/P-P stripes. First, Pol II elongation may drag “sticky” promoters and enhancers along with the transcription machinery during elongation until colliding with another large protein complex such as a transcription preinitiation complex or cohesin complex. As the elongating complex is unable to pass through these structural blockades, we often observe “dots” at the ends or intersections of stripes. Our finding that acute Pol II transcription inhibition disrupts both enhancer and promoter stripes is consistent with this model. Second, when the transcription machinery and cohesin complex collide, Pol II can assist the cohesin-associated loop extrusion machinery (*44*), which may also result in the formation of E-P/P-P stripes. The recent finding that CTCF/cohesin-mediated stripes associate with active enhancer and gene expression at the *Epha4* developmental locus provides an example of this model with a functional readout (*45*). However, yeast have a similar transcription machinery and SMC complex, but no stripe and loop structures were found during log phase, implying that active replication may act as a dominant force to disrupt these transcription-related structures in yeast (*22, 23*). Third, multiple promoters and enhancers may be trapped within a “regulatory protein/DNA hub” (fig. S8D). These interactions are expected to be sufficiently stable and in close enough proximity to be captured by chemical crosslinking. Evidence from pile-up analyses of enhancer factors (Nanog, P300) as well as single loci with a stripe linking multiple enhancers supports this mechanism (Fig. 4). Consistently, HiChIP analysis of Klf4 and H3K27ac revealed enhancer hubs that were correlated with cell-type specific gene regulation (*46*). We anticipate that these three models may not be mutually exclusive and could function together *in vivo*. Further functional studies will be required to dissect the detailed mechanisms of E-P/P-P stripe formation.

Gene-specific and local nucleosome folding have also been poorly explored by genome-wide approaches in mammals. We found that, although gene folding is dependent on transcription activity as seen in yeast (*22*), remarkably, an inverse relationship was found in mammals. Unlike the yeast situation, in mammalian cells, gene stripes and intra-gene interactions are highly enriched in actively transcribed genes. This may reflect a more complicated transcriptional regulatory mechanism in mammals. Thus, we envision two potential models to describe the transcription/Pol II-dependent gene folding. First, gene folding may be mediated by “sticky” Pol II as described above (fig. S10D, (i)). Alternatively, the promoter and related cis-regulatory elements could come together inside a Pol II hub, while Pol II and other factors deliver a range of chromatin elements into the hub for transcription (fig. S10D, (iI)). Acute Pol II inhibition only disrupts gene and enhancer stripes, but global chromatin folding remained unchanged, indicating that preinitiation complex assembly and Pol II elongation might only account for a portion of chromatin folding, particularly at the scale of individual genes. Nevertheless, we cannot rule out the possibility that a short period of Pol II inhibition may not completely shut down transcriptional activity and evict preloaded Pol II that could continue elongating through a long gene. Furthermore, although acute depletion of CTCF and cohesin largely abolishes TADs, how they may affect microTADs and gene-level folding remains to be determined. High-resolution Micro-C maps can be used to tackle these questions and provide additional links between fine-scale chromatin folding (E-P/P-P stripes) and gene regulation. Here we have revealed a previously unknown layer of chromatin folding in mammals. In the future, we expect that further studies that combine nucleosome-resolution chromatin maps with live-cell single-molecule imaging or single-cell technologies will further refine our understanding of chromatin folding and its function during mammalian gene regulation.

## ACKNOWLEDGMENTS

We thank David McSwiggen, Claire Darzacq, Yu Chen, and other members of the Tjian and Darzacq labs for comments on the manuscript. Nil Krietenstein discussed the Micro-C experiments. Peter Sudmant discussed the boundary analysis. Gouhong Li shared the raw data of the two-start zigzag model. Sebastian Balaciu illustrated the 30 nm chromatin models.

### Fundings

This work was supported by NIH grants UO1-EB021236 and U54-DK107980 (XD), the California Institute of Regenerative Medicine grant LA1-08013 (XD), Koret UC Berkeley-Tel Aviv University Initiative grant (TSH&XD), and by the Howard Hughes Medical Institute (003061, RT). TSH is a postdoc fellow of the Koret UC Berkeley-Tel Aviv University Initiative. ES is an undergraduate fellow of SURF L&S at UC Berkeley. ASH was a postdoctoral fellow of the Siebel Stem Cell Institute and is supported by a NIH K99 Pathway to Independence Award (NIGMS K99GM130896). This work used the Vincent J. Coates Genomics Sequencing Laboratory at UC Berkeley, supported by NIH S10 Instrumentation Grants 10RR029668 and S10RR027303.

### Author contributions

TSH, ES, ASH, CC, RT, and XD designed and conceived of the project. TSH lead and performed experiments and data analysis. ES performed chromatin dot analysis. ASH and CC generated cell lines and contributed to data analysis. TSH drafted the manuscript and all authors edited the manuscript. XD and RT supervised the project.

### Competing interests

No competing interests.

### Data availability

All data are available at GSE130275.

## SUPPLEMENTARY MATERIALS

Materials and Methods

Figures S1-S12

Table S1

References

