## Supplemental Materials for "Resolving the 3D landscape of transcription-linked mammalian chromatin folding"

#### Supplementary Materials for
#### Resolving the 3D landscape of transcription-linked mammalian chromatin folding

Tsung-Han S. Hsieh[1,2,3,4,5], Elena Slobodyanyuk[1,2,3,5], Anders S. Hansen[1,2,3,5], Claudia Cattoglio[1,2,3,5], Oliver J. Rando[6], Robert Tjian[1,2,3,5*], Xavier Darzacq[1,2,3,4*]

[1]Department of Molecular and Cell Biology, UC Berkeley
[2]Li Ka Shing Center for Biomedical and Health Sciences
[3]CIRM Center of Excellence
[4]Center for Computational Biology, University of California, Berkeley, Berkeley, United States;
[5]Howard Hughes Medical Institute, University of California, Berkeley, Berkeley, United States;
[6]Department of Biochemistry and Molecular Pharmacology, University of Massachusetts Medical School, Worcester, MA, United States

**This PDF file includes:**

Materials and Methods
- Micro-C experiment
- Data analysis
- Step-by-step Micro-C protocol

Supplemental figures and figure legends
- Figures S1-S12

Supplemental tables
- Table S1

Supplemental references

**Materials and Methods**

Micro-C protocol for mammals was modified from the original protocol for yeast in (*1*, *2*). The protocol was optimized for the input cell number from 1k to 5M and first applied to the mammalian system in (*3*). We first briefly summarize the critical steps and concepts in the Micro-C method, and then provide detailed step-by-step instructions.

**Micro-C experiment**

1. Cell culture and crosslinking

Here, we performed a dual crosslinking protocol to fix protein-DNA and protein-protein interactions. In addition to formaldehyde, we used the non-cleavable and membrane-permeable protein-protein crosslinker DSG (disuccinimidyl glutarate, 7.7Å) or EGS (ethylene glycol bis(succinimidyl succinate), 16.1Å) to crosslink the primary amines between proximal proteins. The dual-crosslinking method significantly increases the signal-to-noise ratio of Micro-C data in yeast (*2*).

In brief, 1k – 5M cells were resuspended by trypsin and fixed by freshly made 1% formaldehyde at room temperature for 10 minutes. The crosslinking reaction was quenched by adding Tris buffer (pH = 7.5) to final 0.75 M at room temperature. Fixed cells were washed twice with 1X PBS and protein-protein interactions fixed by 3 mM DSG for 45 minutes at room temperature. The DSG solution was freshly made at a 300 mM concentration in DMSO and diluted to 3 mM in 1X PBS before use. The crosslinking reaction was quenched by 0.75 M Tris buffer and washed twice with 1X PBS. Crosslinked cells were snap-frozen in liquid nitrogen and stored at -80°C (pellets are stable for up to a year). Note that freshly made crosslinking solution is critical to producing high-reproducibility Micro-C data, and Tris buffer is a faster and stronger quenching agent than glycine.

2. Chromatin fragmentation by Micrococcal nuclease (MNase)

In Hi-C protocol, solubilizing chromatin by SDS allows restriction enzymes to access and cut their target sequences, while SDS-solubilization of chromatin seems to impede nucleosome-resolution mapping for Micro-C. In our preliminary test, SDS detergent appears to "over-solubilize" chromatin – MNase has an equal chance to access nucleosomal DNA and linker DNA, resulting in a smearing nucleosome ladder and an obscure nucleosome occupancy. Consequently, instead of using an anionic/denaturing detergent like SDS we only use the nonionic detergent NP-40 to mildly permeabilize the nuclear membrane. The nonionic detergent condition supposedly retains intact nuclei for "in situ" mapping of chromatin interactions.

To this end, intact nuclei were extracted by treating cells with Micro-C Buffer #1 (50 mM NaCl, 10 mM Tris-HCl pH = 7.5, 5 mM $MgCl_2$, 1M $CaCl_2$, 0.2% NP-40, 1x Protease Inhibitor Cocktail) for 20 minutes on ice. Chromatin in the permeabilized nuclei was digested with a pre-titrated MNase concentration at 37°C for 10 minutes, which generates about 90% of mononucleosomes and 10% of di-nucleosomes (fig. S1A). The ratio has been tested to yield the best signal-to-noise ratio in yeast Micro-C data, retaining long enough DNA ends on mononucleosomes for ligation and reducing unligated products introduced by undigested dimers as well. MNase digestion was stopped by adding 4 mM EGTA and completely inactivated by incubating at 65°C for 10 minutes. Digested chromatin was washed twice with ice-cold Micro-C Buffer #2 (50 mM NaCl, 10 mM Tris-HCl pH = 7.5, 10 mM $MgCl_2$).

3. End repairing and labeling

MNase-digested chromatin exhibits various types of DNA ends, including 5' overhangs, 3' overhangs, and blunt ends. The digested ends cannot be ligated by T4 DNA ligase, as MNase leaves a 3'-phosphate (3'-P) and a 5'-hydroxyl group (5'-OH), rather than 5'-P and 3'-OH on the DNA ends. To generate ends compatible with T4 DNA ligase (blunt ends with 5'-P) for proximity ligation, digested chromatin was subjected to multiple steps of biochemical enzyme reactions: 1) T4 Polynucleotide Kinase catalyzes the addition of 5'-P and removal of 3'-P to generate ligatable ends on nucleosomal DNA. Chromatin was incubated with T4 PNK in Micro-C end-repairing buffer (50 mM NaCl, 10 mM Tris-HCl pH = 7.5, 10 mM $MgCl_2$, 100 ug/mL BSA, 2 mM ATP, 5 mM DTT) at 37 °C for 15 minutes. 2) DNA Polymerase I Klenow Fragment can remove 3' overhangs (3' → 5' exonuclease) or fill-in 5' overhangs (5'→ 3' polymerase) to form blunt ends. In "dNTP-depletion" conditions, Klenow Fragment only acts to remove the 3' overhang and keeps chewing into nucleosome DNA until blocked by the crosslinked histones. Polymerase activity will dominate the exonuclease activity upon addition of dNTPs. We thus employed the dual functions of Klenow Fragment to generate biotin-labeled blunt ends, by incubating chromatin with Klenow Fragments in the Micro-C end-repairing buffer with no dNTPs at 37 °C for 15 minutes. The blunting and labeling reaction was triggered upon adding biotin-dATP, biotin-dCTP, dGTP, and dTTP to a final concentration of 66 mM for each. Incubation for 45 minutes at room temperature is sufficient to convert most MNase-digested ends to blunt ends for proximity ligation. T4 PNK and Klenow Fragment were inactivated by adding 30 mM EDTA and incubating in 65°C for 20 minutes. Biotin-labeled chromatin was washed once by ice-cold Micro-C Buffer #3 (50 mM Tris-HCl pH = 7.5, 10 mM $MgCl_2$). Note that T4 DNA Polymerase also produces a similar result to Klenow Fragment, but a stronger 3' → 5' exonuclease activity makes the reactions harder to control.

4. Proximity ligation and removal of biotin-dNTP from unligated ends

Since the protocol retains intact nuclei throughout the procedure, we found that in situ/in nuclei ligation is very fast and robust, and there is no benefit of an excessive dilution volume or a prolonged ligation time to the signal-to-noise ratio. To obtain optimal results, DNA ends between crosslinked nucleosomes were ligated by T4 DNA Ligase in 500 µl solution at room temperature for at least 2 hours.

Once two proximal nucleosomes are ligated together, the only meaningful biotin signals for later detecting nucleosomal contacts are protected in the middle of ligated di-nucleosomes. Removing biotin-DNA at the ends of chromatin fragments significantly increases signal-to-noise in the Micro-C maps and reduces the ratio of undigested di-nucleosomes in Micro-C data. To this end, biotin-dNTPs on the unligated ends were removed by exonuclease III (3' → 5' exonuclease) at 37 °C for at least 15 minutes of incubation.

5. Micro-C library preparation

To specifically extract the ligated dinucleosomal DNA, the deproteinized chromatin was purified and separated on a low-melting agarose gel. A band at the size of 200-400bp corresponding to the ligated dimers was gel-extracted for library preparation (fig. S1A). The purified DNA with biotin-dNTPs was captured by Dynabeads® MyOne™ Streptavidin C1. Standard library preparation protocol including end-repair, A-tailing, and adapter ligation was performed on beads with the NEBnext Ultra II kit. An optimal PCR cycle for final library amplification was determined by quantification PCR, typically between 5 – 10 cycles. The sequencing library was amplified by Kapa HiFi PCR enzyme with the lowest possible cycles to reduce PCR duplicates (fig. S1A). The library was sequenced by paired-end 50x50 or 100x100 in Illumina HiSeq 4000 sequencer (Vincent J. Coates Genomics Sequencing Laboratory at UC Berkeley, supported by NIH S10 OD018174 Instrumentation Grant). Typically, total valid contacts consist of 9.7% of inter-chromosomal contacts and 90.3% of intra-chromosomal contacts, of which 61.4% are shorter than 20 kb, and 38.6% are longer than 20 kb (fig. S1B).

6. Technical aspects of the Micro-C method

As we discussed before, Micro-C not only recapitulates most chromosome features previously identified by Hi-C, but also captures additional finer-scale chromatin structures below the kb-scale. Here, we highlight some key advantages of using Micro-C to study mammalian chromosome folding. 1) Micro-C measures interaction between nucleosomes – the basic unit of chromatin, instead of between uneven chunks of chromatin fragmented by restriction enzymes. This modification essentially increases the mapping resolution to single-nucleosome resolution and enables the study of chromatin folding at a scale below kilobase. 2) The Micro-C protocol uses a non-cleavable crosslinker to fix protein-protein interactions. Studies have reported that formaldehyde crosslinking is usually slow and incomplete, and tends to be reversed with time. Proteins can still freely diffuse up to hours after formaldehyde treatment (*4*). The uncertainty of crosslinking quality inevitably results in a poor resolution, especially in highly dynamic chromatin regions. To examine the effect of crosslinking on detecting chromatin structures, we performed Micro-C with a time course of crosslinking from 1 to 15 minutes (fig. S12). As we expected, relatively stable chromatin structures like TADs can be identified within 5 minutes of crosslinking, while loops become clearer after 5-10 minutes of crosslinking. Undoubtedly, these measurements can provide only a clue for optimizing the crosslinking conditions and a rough picture of chromatin dynamics. To gain a real-time perspective of chromatin dynamics, live single-molecule tracking of TADs and loops will be necessary to obtain a solid conclusion. 3) Micro-C has a higher sensitivity to detect chromatin loops. Hi-C typically requires over 1 billion reads to visualize a single loop structure. Surprisingly, we can start observing many loops with ~50-100M reads by Micro-C (fig. S10A), suggesting that the method provides a cost-effective option for loop-centered studies. 4) Studying the fine-scale chromatin folding such as gene folding, E-P/P-P links, and 30 nm chromatin structure requires nucleosome-resolution chromatin maps. These structures are hardly detectable at the genome-wide scale by other 3C-based techniques. 5) The Micro-C protocol also works with low-input cell numbers, as 1k – 10k sorted cells have been successful in our tests. We envision that single-cell Micro-C can be developed with the recent advances in single-cell technologies.

**Data analysis**
All FASTQ, Valid pairs, COOL, and HIC files are publicly available at GSE130275.

1. Mapping and pairing Micro-C contacts

Valid Micro-C contact read pairs were obtained from the HiC-Pro analysis pipeline (*5*). The detailed description and code can be found at https://github.com/nservant/HiC-Pro. Fastq files were mapped to mouse mm10 genome by Bowtie2 with 'very sensitive' mode. Aligned reads were paired. Pairs with multiple hits, low MAPQ, singleton, dangling end, self-circle, and PCR duplicates were removed. Output files containing all valid pairs were used in downstream analysis.

2. Binning and data normalization for Micro-C maps

We generated 100bp-bin files of mouse mm10 genome for assigning Micro-C contact pairs, which virtually resembles the "nucleosome" resolution. Tab-delimited valid pairs were mapped to the 'pseudo' nucleosome occupancy files and converted to HDF5 format as .COOL files using the COOLER package (https://github.com/mirnylab/cooler) (*6*), or converted to .HIC files using the JUICER package (https://github.com/aidenlab/juicer) (*7*). Regions with low mappability and high noise were precluded before matrix normalization. Contact matrices were then balanced by using iterative correction (IC) for COOL files (*8*) or Knight-Ruiz (KR) (*9*) for HIC files. We assume that the nucleosome occupancy of Micro-C maps should be corrected with matrix balancing. Both normalization methods produce visually equal quality of contact maps.

3. Browsing Micro-C contact maps

Processed Micro-C data can be converted to the standard 4DN formats such as .COOL and .HIC, with multiple resolutions from 100 bp to Mb. A compilation of multiple resolutions of .COOL (.mCOOL) can be visualized on the HiGlass browser (http://higlass.io) (*10*), and .HIC files are compatible with the Juicebox browser (https://github.com/aidenlab/Juicebox) (*11*). All processed files can be found at GSE130275. In this study, all browser snapshots of Micro-C/Hi-C contact matrices and the 1D browser tracks (e.g., ChIP-seq, ATAC-seq, MNase-seq) were generated by the HiGlass browser unless otherwise mentioned.

4. Reproducibility test

Reproducibility of Micro-C data was evaluated by three algorithms independently (fig. S1D-F). The packages can be found at https://github.com/kundajelab/3DChromatin_ReplicateQC. QuASAR calculates the correlation of values in two distance-based transformed matrices (https://github.com/bxlab/hifive) (*12*). GenomeDISCO measures the difference in two graph diffusion smoothed contact maps (https://github.com/kundajelab/genomedisco) (*13*). Hi-Rep calculates reproducibility by a weighted sum of correlation coefficients (https://github.com/qunhualilab/hicrep) (*14*).

5. Contact decaying curve analysis

We only used intra-chromosomal contact pairs to calculate the contact probability in bins with exponentially increasing widths from 200 bp to 10 Mb, or with single base pair from 200 to 2000 bp. Contacts with a distance shorter than 200 bp were removed from the analysis to minimize potential noise introduced by self-ligation or undigested DNA products. Decaying curves in this study were normalized to the total number of contact pairs. The orientations of ligated DNA are noted as “IN-IN (+/-),” “IN-OUT (+/+),” “OUT-IN (-/-),” and “OUT-OUT (-/+)” according to the readouts of Illumina sequencing (*1*). “UNI” pairs are the combination of “IN-OUT” and “OUT-IN” because both orientations are theoretically interchangeable. Schematics illustrating the corresponding orientation of nucleosome interactions are shown in fig. S11C. In Fig. 1B, we only showed the decaying curves of the “UNI” pairs for clarity.

6. Chromosome compartment analysis

Chromosome compartments were identified by Principal Component Analysis (PCA) of the contact matrix at 100-kb resolution (fig. S12C). The eigenvectors of the first component typically represent the compartment profile in Hi-C data (*15*), as positive values are the A compartment (gene-rich/active chromatin) and negative values are the B compartment (gene-poor/inactive chromatin). The saddle plot shown in fig. S10A represents the rearrangement and aggregation of genome-wide distance-normalized contact matrix with the order of increasing eigenvector values.

7. Chromatin domain analysis

We used insulation score analysis (*16*) to identify sharp changes of chromatin interactions, which typically represent the domain boundaries (Fig. 2 and fig. S4). To identify the fine-scale chromatin structure, we analyzed insulation profiles with Micro-C contact matrices at 200-bp, 400-bp, 600-bp, 800-bp, 1-kb, 2-kb, 4-kb, 10-kb, and 20-kb resolutions. We used sliding windows 10 times larger than the given resolution, e.g., a 2000-bp sliding window for 200-bp resolution. Similar results are obtained with a 20- or 100-times larger sliding windows. The signal within the sliding window was assigned to the corresponding bin across the entire genome. The insulation scores were normalized to the log2 ratio of the individual score and the mean of the genome-wide averaged insulation score. Chromatin boundaries can be identified by finding the local minima along with the normalized insulation score. Boundaries overlapping with low mappability regions were removed from the downstream analysis. For aggregate domain analysis (“ADA”) in fig. S10A, each domain was rescaled to a pseudo-size by $N_{i,j}$=(($C_i$-$D_{start}$)/($D_{end}$-$D_{start}$), ($C_j$-$D_{start}$)/$D_{end}$-$D_{start}$)), where $C_{i,j}$ is a pair of contact loci within domain D that is flanked by $D_{start}$ and $D_{end}$, and $N_{i,j}$ is a pair of the rescaled coordinates. The rescaled domains can be aggregated at the center of the plot with ICE or distance normalization. This analysis also applied to the rescaled pile-up analysis of gene structure in fig. S9A-B.

8. Chromatin loop/dot analysis

Chromatin loops in this study were identified by using the HiCCUPS algorithm (*17*), which is available in the JUICER package. Loops were called at multiple resolutions (0.5, 1, 5, 10 kb) of KR-normalized Micro-C contact matrices, and filtered by a false discovery rate at 0.1. After merging loops called by multiple resolutions, we identified 29,548 loops with 2.64B reads of Micro-C dataset and 14,372 loops with 1.3B reads of data, but only 6,005 loops with 3.3B reads of Hi-C data (*18*). We noticed that current loop callers yield undesirably high false positive and negative loop identifications, and the calling accuracy heavily depends on 1) sensitivity of the method, 2) resolutions and parameters (e.g., peak and window size) used to call loops, and 3) sequencing coverage. For example, we tested multiple resolutions and parameters to identify loops with 2.64B of Micro-C data:

| Resolution (bp) | FDR | Peak size | Window size | Distance to merge | Loop number |
|---|---|---|---|---|---|
| 1000 | 0.1 | 4000 | 10000 | 2500 | **16525** |
| 2500 | 0.1 | 10000 | 20000 | 5000 | **29492** |
| 5000 | 0.1 | 15000 | 30000 | 10000 | **26649** |
| 10000 | 0.1 | 20000 | 40000 | 20000 | **18086** |

We had difficulties determining true positives, as five proximal 1000-bp loops sometimes are identified as a single 5000-bp loop, and 5000-bp loops sometimes are melted to widely spread interactions off the diagonal when zooming into a higher resolution. To avoid these issues, we decided to focus on less-biased, higher resolution, and more robust insulation analysis in this study. In some cases, we used our proofread loop list (by eyes) for specific applications.

Genome-wide loop intensity was assessed by aggregate peak analysis (“APA”) (Fig. 4D-E, 5C and fig. S8A-B, S9D-E, S10A, S12C). Loops were piled up on the center of a 25-kb x 25-kb matrix with a 1-kb resolution of KR-normalized data. Loops within 55 kb of the diagonal were excluded to avoid distance decay effects. The ratio of loop enrichment was calculated by dividing observed contact in a searching window by the expected bottom-left submatrix.

For the target-centered loop analysis, ChIP-seq peaks/transcription start sites (TSSs)/transcription termination sites (TTSs) at distances shorter than 1 Mb were paired (since the majority of loops are formed within a 1-Mb range). We then calculated the local pixel enrichment (as described above) in a searching window for quantification of peak-mediated loops. Pairs with no enrichment or FDR > 0.1 were removed from the analysis. Genome-wide evaluation of target-centered loop formation was assessed by APA.

9. Pile-up analysis

The principle of pile-up analysis is similar to the APA analysis described above. We used targets of interest (e.g., ChIP-seq peaks) as bait to extract either 20-kb (for nucleosome-resolution map) or 400-kb (for 5-kb resolution map) snippets of contact maps from Micro-C data. The coordinate of the target was centered at each snippet. All corresponding snippets were then piled-up together and normalized by the expected matrix (Fig. 4F-G, Fig. 5B and G, and fig. S8C).

10. ChIP-seq analysis

We reanalyzed the 48 public available datasets (Table S1) with the HOMER package (http://homer.ucsd.edu/homer/) (*19*). Peaks were called by the peak analysis function in the package or by MACS2 independently.

11. Boundary prediction

Boundary location and genome-wide data were converted to a binary format by ChromHMM (http://compbio.mit.edu/ChromHMM/) (*20*), where the bin with a boundary and peak is one, and all others are zero. We then trained the program with a series of predictors to predict the boundary location. Redundant factors were removed by the cutoff of the 75$^{th}$ lambda value in Lasso regularization analysis. The strongest predictive factors of boundary location were identified by generalized linear regression analysis. We used the same strategy to find the predictive factors of boundary strength, but using the ChIP-seq single enrichment as the vector rather than the binarized data (Fig. 3B-C and fig. S5E-F).

12. Boundary classification

To dissect the properties of a single microTAD boundary, we first quantified the signal enrichment of 48 genomic datasets at each boundary (Table S1) and then built a covariance matrix for Principal Component Analysis (PCA). Overall, the first three components, which included factors related to transcription and chromatin functions, explained ~77.71% of the total variation (fig. S6A). To classify boundaries into subgroups, we used dimension-reduced data from PCA as input (~90% of the total variance explained in PC1-10) for t-Distributed Stochastic Neighbor Embedding (t-SNE) analysis. Individual boundaries were plotted as embedded points in 2D space, and then color-coded by signal enrichment of the 48 interrogated datasets (Fig. 3F-G and fig. S6B)

13. Stripe decaying curve analysis

We first extracted the chromatin interactions of each “horizontal stripe” and “vertical stripe” that colocalize with the protein-binding sites at 1-kb width, and extend up to 100 Mb away. For example, if there are ~80,000 CTCF peaks, we will get the ~80,000 horizontal stripes and vertical stripes that extend from each CTCF binding sites. The size of each stripe is 1 kb x 100 Mb. We then calculated the interaction decaying behaviors of these stripes with distance.

**Step-by-step Micro-C protocol**

I. Prepare crosslinked chromatin from cell culture.

1. Culture cells in recommended condition.
2. Harvest cells by centrifugation for 5 min at 800xg at room temperature.
3. Resuspend cells in base media (w/o FBS) in a concentration of $1x10^6$ cells/mL (max. 30mL in 50mL tube).
4. Add 16% Formaldehyde to a final concentration of 1% (e.g. 2mL for 30mL sample). Incubate for 10 min at room temperature with mixing.
5. Add 2M Tris pH 7.5 to a final concentration of 0.75 M to quench the reaction (e.g. 18.75 mL for 30 mL sample).
   Incubate for 5 min at room temperature.
   Centrifuge for 5 min at >800xg at room temperature. Discard supernatant.
6. Wash cells twice by 1X PBS in a concentration of $1x10^6$ cells/mL.
   Centrifuge for 5 min at >800xg at room temperature. Discard supernatant.
7. Freshly prepare long crosslinkers solution as described in the table:

| Crosslinkers | MW | Spacer (Å) | Stock | Working |
|---|---|---|---|---|
| DSG | 326.26 | 7.7 | 300 mM in DMSO | 3.0 mM in PBS |
| EGS | 456.36 | 16.1 | 300 mM in DMSO | 3.0 mM in PBS |

8. Resuspend cell pellet in long crosslinker solution in a concentration of $1x10^6$ cells/mL. Incubate for 45 min at room temperature with mixing.
9. Add 2 M Tris pH 7.5 to a final concentration of 0.75 M to quench the reaction.
   Incubate for 5 min at room temperature.
   Centrifuge for 5 min at >800xg at room temperature. Discard supernatant.

10. Wash cells twice by 1X PBS in a concentration of $1x10^6$ cells/mL.
Centrifuge for 5 min at >800xg at room temperature. Discard supernatant.
11. Snap freeze cell pellets by liquid nitrogen.

<u>II. Digest crosslinked chromatin by micrococcal nuclease</u>

1. Fresh, complete MB#1:

| Total | 10 mL | 5 mL | 2.5 mL | Final |
|---|---|---|---|---|
| MB#1 | 9700 µL | 4850 µL | 2425 µL | 50 mM NaCl, 10 mM Tris, 5 mM $MgCl_2$, 1 mM $CaCl_2$, 0.2% NP-40, 1X PIC |
| 10% NP-40 | 200 µL | 100 µL | 50 µL | |
| 100X proteinase inhibitors | 100 µL | 50 µL | 25 µL | |

2. Resuspend cell pellet in complete MB#1 in a concentration of $1x10^6$ cells/100µL.
Incubate for 20 min on ice.
Centrifuge for 5 min at ~10000xg for at 4°C. Discard supernatant.
3. Wash nuclei pellet in complete MB#1 in a concentration of $1x10^6$ cells/100µL.
Centrifuge for 5 min at ~10000xg for at 4°C. Discard supernatant.
4. Resuspend nuclei pellet in complete MB#1 in a concentration of $1x10^6$ cells/100µL.
5. Add the appropriate amount of MNase to digest chromatin to 90% Monomer/10% Dimer.
Incubate for 10 min at 37°C with shaking at 850 rpm.
6. Add 500 mM EGTA in a final concentration at 4 mM to stop the reaction.
Incubate for 10 min at 65°C.
Centrifuge for 5 min at >10,000xg at 4°C. Discard supernatant.
7. Wash/Rinse nuclei pellet in 1 mL of cold MB#2 twice.
Centrifuge for 5 min at >10,000xg at 4°C. Discard supernatant.

<u>III. Repair fragment ends</u>

| A. END-CHEWING | | | |
|---|---|---|---|
| **Total** | **100 µL** | **50 µL** | **Final** |
| Chromatin | pellet | pellet | 50 mM NaCl, 10 mM Tris, 10 mM MgCl2, 100 µg/mL BSA<br>2 mM ATP<br>5 mM DTT |
| 10X NEBuffer 2.1 | 10 | 5 | |
| 100 mM ATP | 2 | 1 | |
| 100 mM DTT | 5 | 2.5 | |
| H2O | 68 | 34 | |
| 10 U/µL T4 PNK | 5 | 2.5 | ~10 U/1µg DNA |
| → Incubate for 15 min at 37°C. | | | |
| 5 U/µL Klenow Fragment | 10 | 5 | ~10 U/1µg DNA |
| → Incubate for 15 min at 37°C. | | | |

| B. END-LABELING | | | |
|---|---|---|---|
| **Total** | **150 µL** | **75 µL** | **Final** |
| Chromatin | 100 µL | 100 µL | 66 mM dNTP / each<br>33 mM NaCl, 23 mM Tris, 10 mM MgCl2, 100 µg/mL BSA<br>1.67 mM ATP<br>6.67 mM DTT |
| 1 mM Biotin-dATP | 10 | 5 | |
| 1 mM Biotin-dCTP | 10 | 5 | |
| 10 mM dTTP + dGTP | 1 | 0.5 | |
| 10X T4 DNA Ligase Buffer | 5 | 2.5 | |
| 20 mg/mL BSA (200X) | 0.25 | 0.125 | |
| H2O | 23.75 | 11.875 | |
| → Incubate for 45 min at 25°C with interval mixing.<br>→ Add 500 mM EDTA to 30 mM final concentration → 65°C for 20 min.<br>→ Centrifuge for 5 min at > 10,000xg at 4°C. Discard supernatant. | | | |

| |
|---|
| → Rinse once with 1 mL of cold MB#3.<br>→ Centrifuge for 5 min at > 10,000xg at 4°C. Discard supernatant. |

IV. Proximity ligation & purge of unligated ends

| **A. LIGATION** | | | |
|---|---|---|---|
| **Total** | **500 µL** | **250 µL** | **Final** |
| Chromatin | pellet | pellet | 50 mM Tris, 10 mM $MgCl_2$, 1 mM ATP, 10 mM DTT<br>100 µg/mL BSA |
| Water | 422.5 | 211.25 | |
| 10X T4 DNA ligase buffer *w/ ATP* | 50 | 25 | |
| 20 mg/mL BSA (200X) | 2.5 | 1.25 | |
| 400 U/µL T4 DNA ligase | 25 | 12.5 | |
| → Incubate for > 2.5 hours at room temperature with slow rotation.<br>→ Centrifuge for 5 min at >16000xg at 4°C. Discard supernatant. | | | |

| **B. REMOVE BIOTIN-dNTP FROM UNLIGATED ENDS** | | | |
|---|---|---|---|
| **Total** | **200 µL** | **100 µL** | **Final** |
| Chromatin | pellet | pellet | 10 mM Bis-Tris-Propane-HCl, 10 mM $MgCl_2$, 1 mM DTT |
| 10X NEBuffer#1 | 20 | 10 | |
| Water | 170 | 85 | |
| 100 U/µL Exonuclease III | 10 | 5 | |
| → Incubate at 37°C for 15 min with interval mixing. | | | |

| **C. REVERSE CROSSLINKING** | | | |
|---|---|---|---|
| **Total** | **250 µL** | **125 µL** | **Final** |
| Chromatin | 200 µL | 100 µL | 2 mg/mL Proteinase K = 2X<br>1X SDS |
| 80X Proteinase K | 25 | 12.5 | |
| 10% SDS solution | 25 | 12.5 | |
| → Incubate at 65°C for overnight. | | | |

V. Di-nucleosomal DNA purification

1. **Phenol:Chloroform:Isoamylalcohol (PCI) extraction:**
   → Add 1X volume of PCI → Vortex for 20 sec → Spin for 10min at 19800xg at room temperature → Keep upper layer.
2. **Purify DNA by ethanol precipitation:**
   → Add 0.1X volume of sodium acetate and 2.5X volume of 100% ethanol → Incubate for >1 hr at -80°C → Spin for 15min at 19800xg at 4°C → Wash pellet by 75% ethanol → Spin for 5 min at 19800xg at 4°C → Air dry pellet for 10min at room temperature → Resuspend pellet in 50 µL of TE buffer (+1X RNase A) → Incubate for >30min at 37°C → ZymoClean.
3. **Size-selection for di-nucleosomal DNA:**
   → Separate Monomer and Dimer by running on 3.5% TAE / 3% TBE NuSieve agarose gel → Cut band at size 250–400 bp (Avoid cutting < 200 bp to eliminate monomer) → ZymoGel purification and resuspend DNA in 18 µL of elution buffer → Quantify DNA by Qubit.

VI. Library preparation

| **1. STREPTAVIDIN PURIFICATION** |
|---|
| a) Wash 2.5 µL beads/sample by 1X TBW.<br>b) Resuspend in 150 µL of 2X BW. |

| |
|---|
| c) Mix with 150 µL of DNA sample for 20 min at room temperature.<br>d) Wash twice by 1X TBW @ 55°C w/ interval mixing.<br>e) Wash by 10 mM Tris.<br>f) Resuspend in 25 µL of EB buffer. |

| **2. END REPAIR** | |
|---|---|
| Total | 30 µL |
| Input DNA | 25 |
| End Prep Reaction Buffer | 3.5 |
| End Prep Enzyme Mix | 1.5 |
| Lid heated ≥ 75°C<br>→ 30 min @ 20°C w/ interval mixing<br>→ 30 min @ 65°C<br>→ Hold @ 4°C | |

| **3. ADAPTER LIGATION** | |
|---|---|
| Total | 46 µL |
| Input DNA | 30 |
| Adapter for Illumina | 0.5 |
| Ligation Master Mix | 15 |
| Ligation Enhancer | 0.5 |
| → 30 min @ 20°C w/ interval mixing | |
| USER enzyme | 1.5 |
| → 15 min @ 37°C w/ interval mixing | |

| **4. BEAD WASH** |
|---|
| a) Wash by 1X TBW @ 55°C w/ interval mixing.<br>b) Wash by 10mM Tris.<br>c) Resuspend in 20 µL of EB buffer. |

| **5. MINIMUM PCR & PURIFICATION** | |
|---|---|
| Total | 10 µL |
| Streptavidin beads | 1 |
| Water | 3 |
| 2X KAPA HiFi Hot Start Mix | 5 |
| 10 µM PE1.0 primer | 0.5 |
| 10 µM PE2.0 primer | 0.5 |

| | | |
|---|---|---|
| Denaturation | 98°C | 45 sec |
| **5-10 cycles** | 98°C | 15 sec |
| | 60°C | 30 sec |
| | 72°C | 30 sec |
| Extension | 72°C | 1 min |
| | 4°C | Hold |
| → Check library size by agarose gel | | |

→ 0.9X Ampure XP beads purification
→ Quantify library by Qubit

VII. Deep sequencing by Illumina PE-50 or PE-100

VIII. Materials

- **DSG (disuccinimidyl glutarate)** (ThermoFisher #20593)
- **EGS (ethylene glycol bis(succinimidyl succinate))** (ThermoFisher #21565)
- **16% Formaldehyde** (Fisher scientific #NC1040701)
- **2 M Tris pH = 7.5** (Sigma Aldrich #)
- **MBuffer#1:** 50 mM NaCl, 10 mM Tris-HCl pH 7.5, 5 mM $MgCl_2$, 1 mM $CaCl_2$, , 0.2% NP-40, 1X PIC.
- **MBuffer#2:** 50 mM NaCl, 10 mM Tris-HCl pH 7.5, 10 mM MgCl2
- **MBuffer#3:** 50 mM Tris-HCl pH7.5, 10 mM MgCl2
- **Micrococcal Nuclease** (Worthington Biochem #LS004798)
  Resuspended from lyophilized powder at 20 U/μl in Tris pH 7.4. Aliquot into tubes upon first use and freeze at –80°C.
- **0.5 M EGTA** (Fisher scientific #)
- **0.5 M EDTA** (Fisher scientific #)
- **T4 DNA Polymerase** (New England Biolabs #M0203)
- **DNA Polymerase I, Large (Klenow) Fragment** (New England Biolabs #M0210)
- **T4 Polynucleotide Kinase** (New England Biolabs #M0201)
- **T4 DNA Ligase** (New England Biolabs #M0202)
- **Exonuclease III (*E. coli*)** (New England Biolabs #M0206)
- **Biotin-14-dATP** (Jena Bioscience #NU-835-BIO14)
- **Biotin-11-dCTP** (Jena Bioscience #NU-809-BIOX)
- **20X Proteinase K solution** (Sigma Aldrich # 3115879001)
  TE with 20 mg/ml proteinase K and 50% glycerol
- **Elution buffer:** 10 mM Tris-HCl pH 7.6
- **TE buffer:** 10 mM Tris-HCl pH 8.0, 1 mM EDTA
- **End-It DNA End-Repair Kit** (EpiCentre BioTechnologies # ER81050)
- **Exo-Minus Klenow DNA Polymerase** (EpiCentre BioTechnologies # KL111)
- **Fast Link DNA Ligation Kit** (EpiCentre BioTechnologies # lk6201)
- **Dynabeads® MyOne Streptavidin C1** (Life Technologies # 65001)
- **KAPA HiFi HotStart ReadyMix** (KAPA Biosystems # KK2601)
- **NEBNext Ultra II** (New England Biolabs #E7645)

#### Supplemental figures and legends

### Supplemental Figure 1

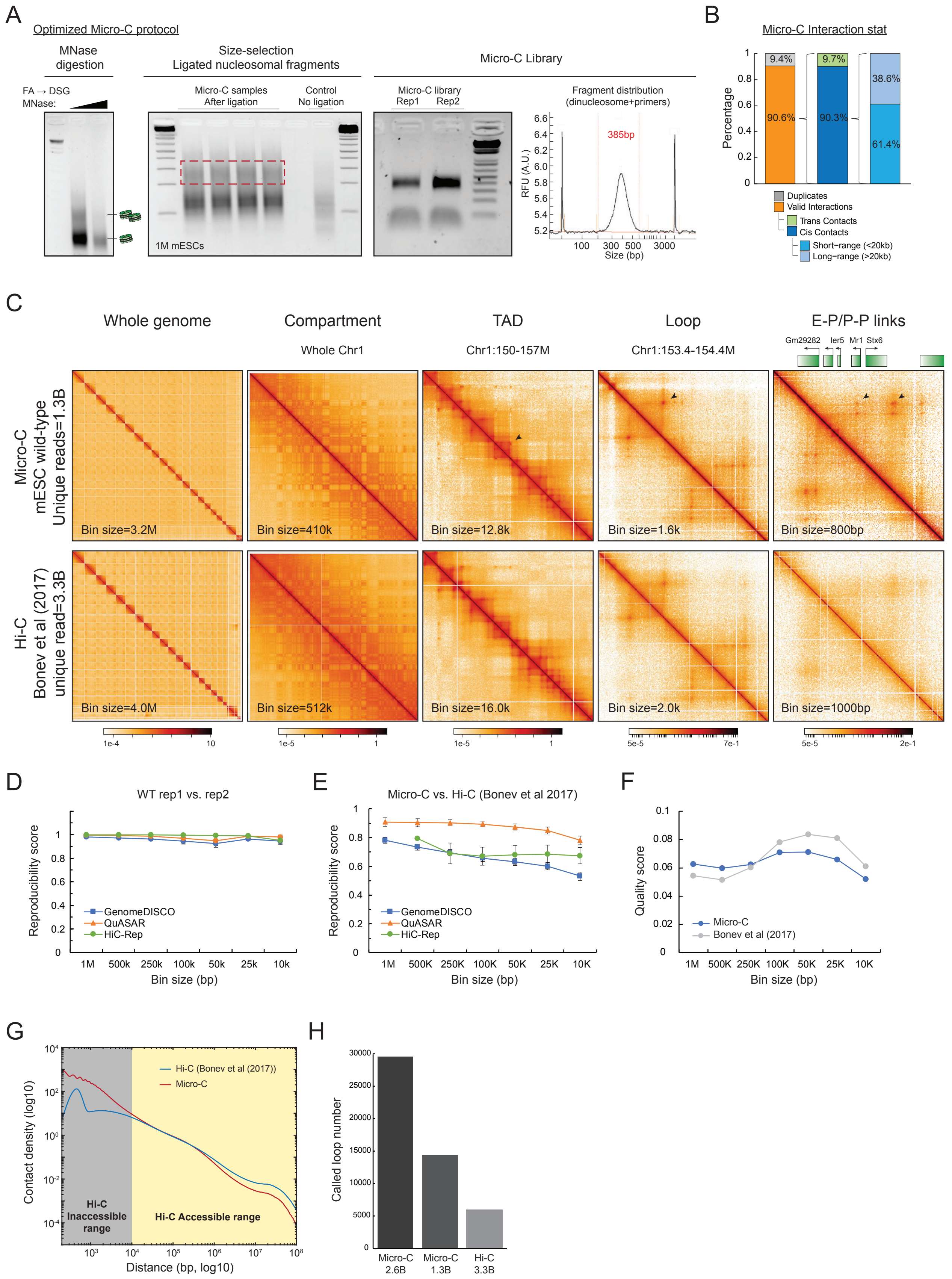

**fig. S1. Micro-C experiment, reproducibility, and quality controls.** (**A**) Example of Micro-C experimental data. The first agarose gel image shows the lower and upper limits for MNase digestion, with slight under-digestion in the first lane and over-digestion in the second lane. Both conditions have been tested to produce highly reproducible results (also see (*1*)), arguing that the level of MNase digestion has a negligible artificial effect on the Micro-C maps as long as the digestion level is within a broad optimal range. The second agarose gel image shows an example of size selection for ligated DNA after proximity ligation, typically between 200 bp to 400 bp. Note that the bands in the Micro-C sample are usually shifted to slightly larger sizes with sharper signals at di-, tri-, and tetramers bands compared to the "No ligation" control, indicating a successful ligation. The third image and the bioanalyzer result show the products of the sequencing library with a sharp band at ~300-500 bp. The optimal PCR cycles should be estimated by quantification PCR, usually ranging between 5 – 10 cycles for 1k – 1M of mESCs. (**B**) Statistics of Micro-C assay for the merged wildtype sample. Typically, the Micro-C library contains over 90% of unique pairs if using a minimal number of PCR cycles. About 90% of valid pairs are cis contacts that consist of 61.4% short-range interactions (<20 kb) and 38.6% long-range interactions (>20 kb), while 10% of pairs are inter-chromosomal contacts. (**C**) Comparison of Micro-C and Hi-C data from Bonev et al. (*18*). Micro-C recapitulates the primary chromatin structures including compartment, TADs, and loops. When zooming into a 350-kb region on chr1, Micro-C shows superior signals to detect chromatin loops, which correspond to E-P/P-P interactions. Black arrows indicate the corresponding chromosome structure in each panel. (**D**) Reproducibility analysis of Micro-C replicates. Reproducibility scores were calculated by three independent algorithms (*21*), including GenomeDISCO (*13*), QuASAR (*12*), and HiC-Rep (*14*). The charts plot reproducibility score (y-axis) at different resolutions (x-axis). All methods reported over 90% of reproducibility rate from 10-kb to 1-Mb resolutions. (**E**) Reproducibility analysis of Micro-C and Hi-C data from Bonev et al. (2017). Reproducibility was calculated by the same algorithms as in (D). Reproducibility scores are 0.8 – 0.9 by QuASAR, 0.6 – 0.8 by GenomeDISCO, and 0.7 – 0.8 by HiC-Rep at 10-kb to 1-Mb resolutions. (**F**) Quality score analysis of Micro-C and Bonev et al. (2017) data. Quality scores were obtained by QuASAR quality control measurement, which is determined by sequencing coverage at a given resolution. Micro-C data in this study has a comparable score to the highest coverage Hi-C data. (**G**) Comparison of interaction decaying rates of Micro-C and Hi-C from Bonev et al. (2017). X-axis is the distance between contact loci from 100-bp to 10-Mb in log10; y-axis is contact density normalized by sequencing depth in log10. Hi-C data loses the resolution in the decaying curve below the distance shorter than 10 kb. (**H**) Comparison of loop calling. Micro-C captures ~5X more loops than Hi-C with similar sequencing depth (Micro-C=2.6B vs. Hi-C=3.3B). The newly-identified loops most likely correspond to the E-P/P-P interactions that were invisible in Hi-C data. However, due to the potential biases of loop calling, we focused on insulation score analysis in the downstream analysis (see also Materials and Methods).

### Supplemental Figure 2

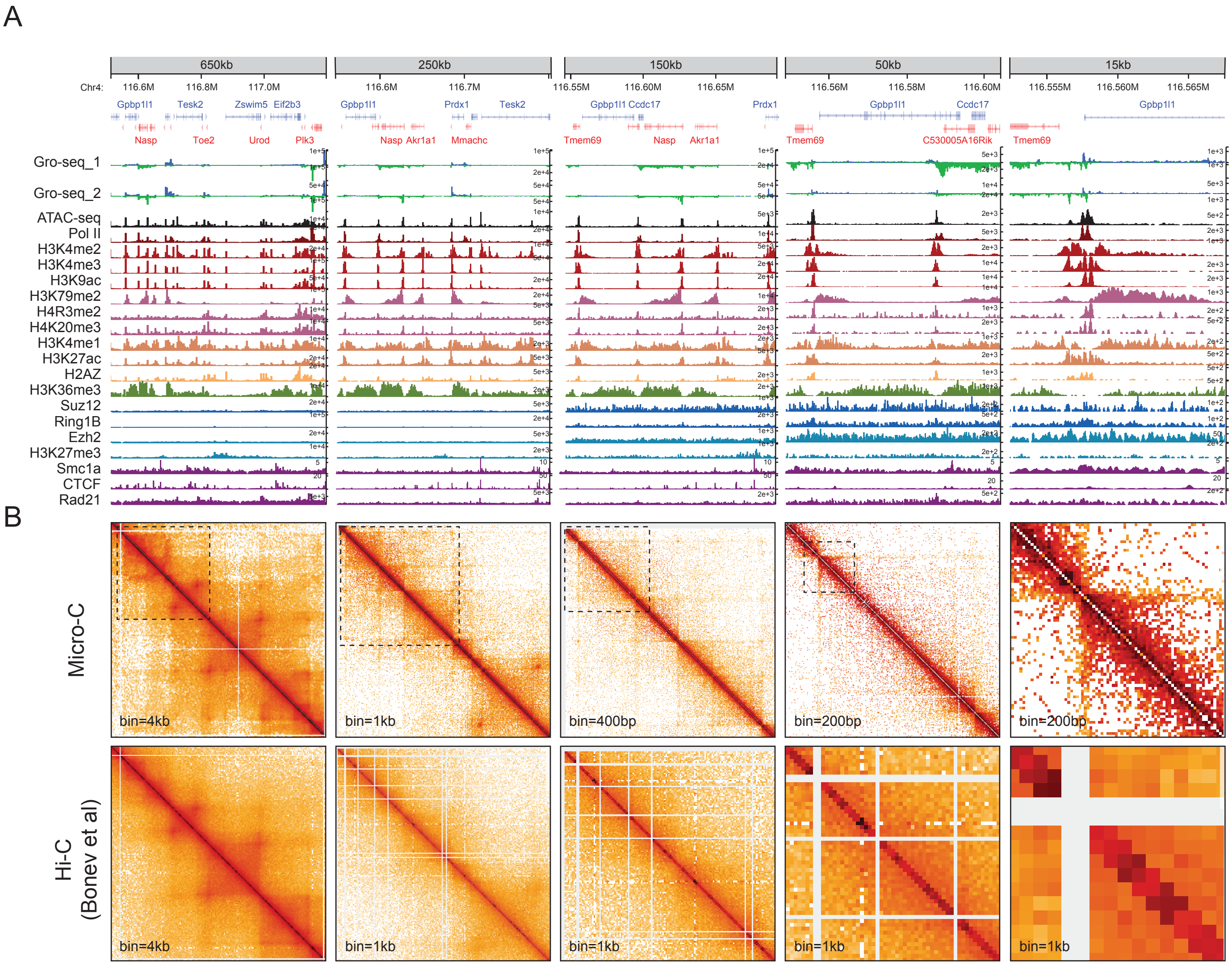

**fig. S2. Additional examples of Micro-C vs. Hi-C contact maps.** Multiple snapshots of 1D chromatin tracks (A) and Micro-C vs. Hi-C contact maps (B) (*18*) on chr4 were sequentially zoomed in from a 650-kb locus to a 15-kb locus. Fine-scale chromatin structures such as short-range self-associating domains (microTADs) appear along the diagonal at 400-bp to 1-kb resolutions, and stripes can be visualized at 200-bp to 400-bp resolutions. MicroTADs are delimited by genomic loci that enrich for nascent RNAs, peaks of ATAC-seq, Pol II, and active histone marks, but independent of CTCF and cohesin peaks. The starting point of the stripe also colocalizes to the active marks. Note that 1 kb is the highest resolution that the Hi-C dataset could reach. Low signal-to-noise and sparse restriction enzyme cutting sites often result in uninterpretable contact maps when zooming in beyond 1-kb resolution.

Supplemental Figure 3

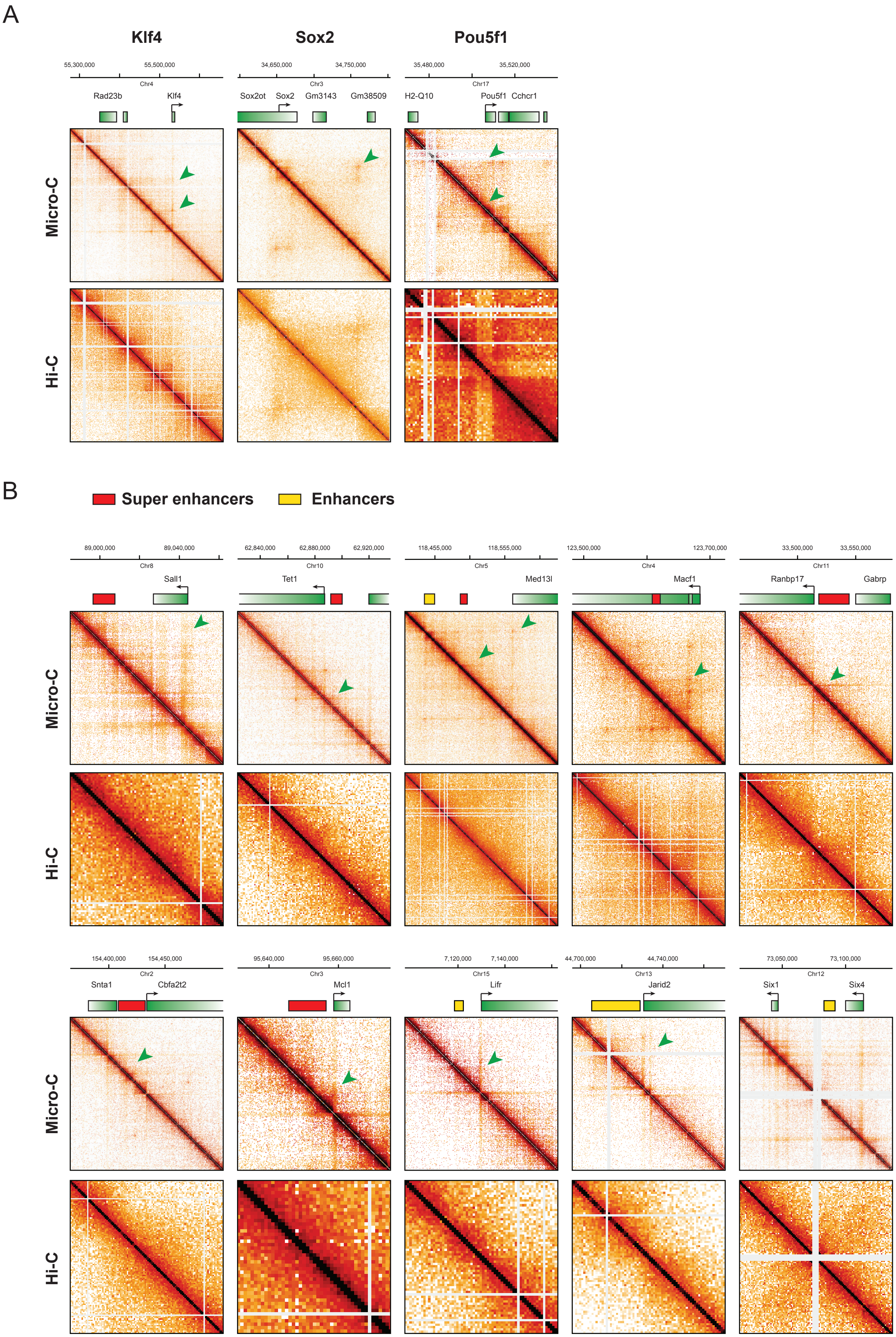

**fig. S3. Micro-C can capture enhancer-promoter structures.** (**A**) Examples of classic enhancer-promoter interactions in mESC. Micro-C analysis robustly identifies E-P links of pluripotency genes *Klf4*, *Sox2*, and *Pou5f1*, but Hi-C (*18*) fails to resolve this level of chromatin structures. (**B**) No obvious structural variations distinguish super-enhancers from regular enhancers. We examined whether super-enhancers and enhancers exhibit structural variations in 3D genome maps. To this end, we selected a list of candidates whose enhancer function has been validated (*22*). Both super-enhancers and enhancers are linked to their promoter by similar architectural stripes, with no significant difference between them in terms of chromatin conformation and interaction intensity.

Supplemental Figure 4

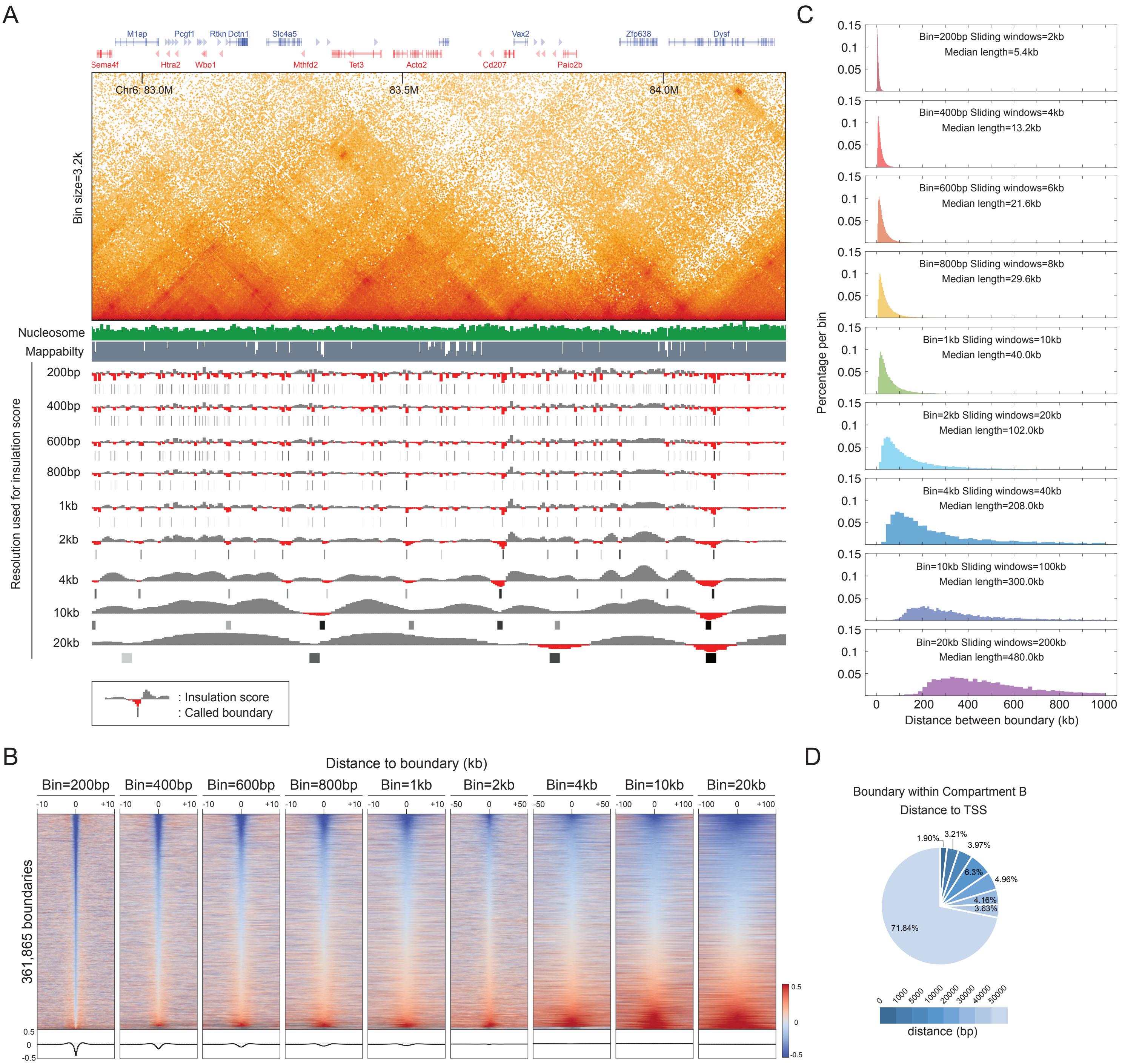

**fig. S4. Boundary identification at multiple resolutions.** (**A**) An example of microTAD boundary identification by the insulation score analysis. Contact map and insulation score tracks show a 1.5-Mb region on chr6. Called boundaries are indicated as black lines in the browser tracks. Map was plotted by a lower resolution (bin size=3.2 kb) of data for visualizing TADs, which were typically characterized by the insulation score analysis at 20 kb (see the bottom panel of the browser tracks). The data also indicates that much finer scale of chromatin domains are hidden within TADs, as the insulation score analyses at 200-800 bp discovered much more chromatin boundaries than the standard 20-kb analysis, stressing the importance of using the nucleosome-resolution map to identify chromatin structures. Nucleosome occupancy and sequencing mappability data shown at the top of the browser track indicate that the majority of the called boundaries are not artifacts resulting from low mappability or nucleosome depletion. Boundaries overlapped with low mappability area were removed from the downstream analysis. (**B**) Additional heatmap and histogram profile of insulation score for Fig. 2B. Heatmaps were plotted by the insulation score analysis at resolutions ranging from 200 bp to 20 kb, and sorted according to the boundary strength in the 200-bp data. Analysis of 20-kb resolution data only recovered 1.2% (n=4384) of the 361,885 microTAD boundaries called by 200-bp data. Genome-wide averaged histograms (below the heatmaps) also show that the strength of the microTAD boundaries gradually decreases from 200-bp to 20-kb resolution. (**C**) Additional examples of the length distribution of microTADs for Fig. 2C. Distribution of the distance between adjacent boundaries is plotted as a histogram across multiple resolutions. X-axis shows the distance between neighboring boundaries, which is equivalent to the domain size identified by the given resolution. (**D**) Distance distribution of the boundaries to the closest TSS in the inactive compartment. Color-coded pie chart shows the percentage distribution of the distance. Over 70% of the microTAD boundary locations in the inactive chromatin are over 50-kb away from TSS, perhaps mediated by transcription-independent mechanisms.

Supplemental Figure 5

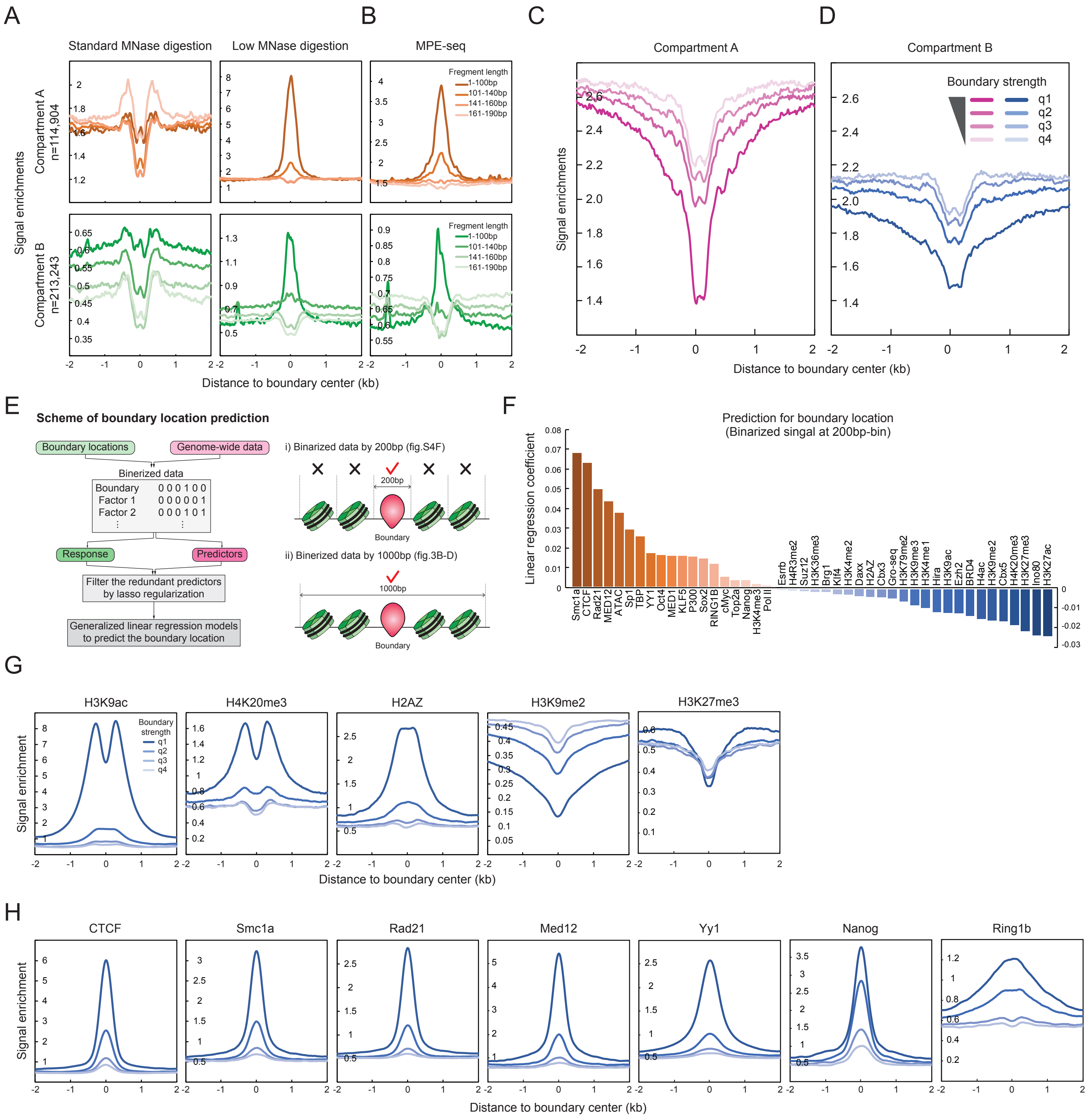

**fig. S5. Features of microTAD boundaries.** (**A-B**) Nucleosome occupancy at microTAD boundaries by MNase-seq and MPE-seq. We applied two independent methods, namely MNase-seq (A) (*24*), and MPE-seq (B) (*25*), to map nucleosome location relative to microTAD boundaries. MNase-seq is the standard method to study nucleosome occupancy, using a standard digestion level to map nucleosome occupancy and an under-digestion condition to map transcription factor binding sites. Typically, nucleosome protection generates 140 – 200-bp fragments, while transcription factors protect fragments smaller than 100 bp. In the standard MNase digestion, the nucleosome signal flanking the microTAD boundaries in active (A) compartments mostly derives from 161-190-bp fragments. Both MNase-seq with a low level of digestion and MPE-seq indicate that the microTAD boundary is enriched at transcription binding sites or cis-regulatory elements, as the 1-100 bp fragment shows the highest signal, in both A and B compartments. (**C-D**) Nucleosome occupancy level at the microTAD boundary. We sorted boundaries to quartiles (q1 to q4) according to their strength and plotted the nucleosome occupancy level (y-axis) ±2-kb around each microTAD boundary quartile (x-axis). Stronger boundaries (q1) correlate with nucleosome depletion levels and localize at nucleosome-depleted regions (NDR). We observed a similar trend in active (**C**) and inactive (**D**) compartments. (**E**) Analysis flow chart for the prediction of boundary location. We first converted the boundary location and genome-wide data to a binary format, as the locus with a boundary or peak is one, and all others are assigned to zero. We then built a series of predictors to train the program to predict the boundary location. The redundant factors were removed by Lasso regularization. We then identified the most predictive factors for boundary location by generalized linear regression analysis. The bin size used for binarizing data affects the prediction results, particularly on the predictive power of histone modifications. As the majority of the microTAD boundaries locate at transcription factor binding sites or fragile nucleosome sites, the 200-bp bin excludes the flanking nucleosomes from the analysis (case i). In Fig. 3B-C, we included nucleosomes for the boundary prediction by using binarized data with 1-kb bin (case ii). (**F**) Prediction of boundary location by the binarized data in 200-bp bins. Y-axis shows linear regression coefficients, and x-axis shows the entry in question. Consistent with our model, transcription factors remain the predictive powers of boundary location, while boundary flanking nucleosomes (histone modifications) were precluded from the prediction at 200-bp resolution. (**G-H**) ChIP-seq signal enrichments at the microTAD boundaries. Histogram curves were plotted by quartile groups of the boundary strength across ±2 kb from the center of the boundary. Overall, the strongest microTAD boundaries are enriched for active marks. Among them, signal enrichments of H3K4me2 in Fig. 3E and Nanog appear to be proportional to the boundary strength, rather than dominated by the first quartile of the signal as in the H3K9ac and H3K4me3 data.

Supplemental Figure 6

A

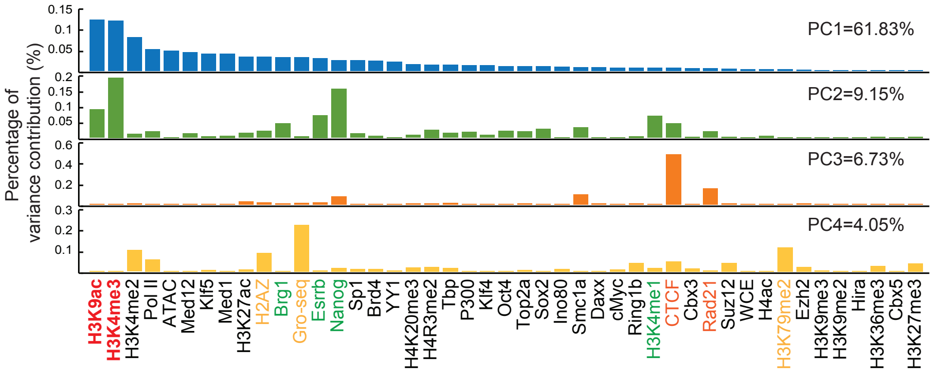

B

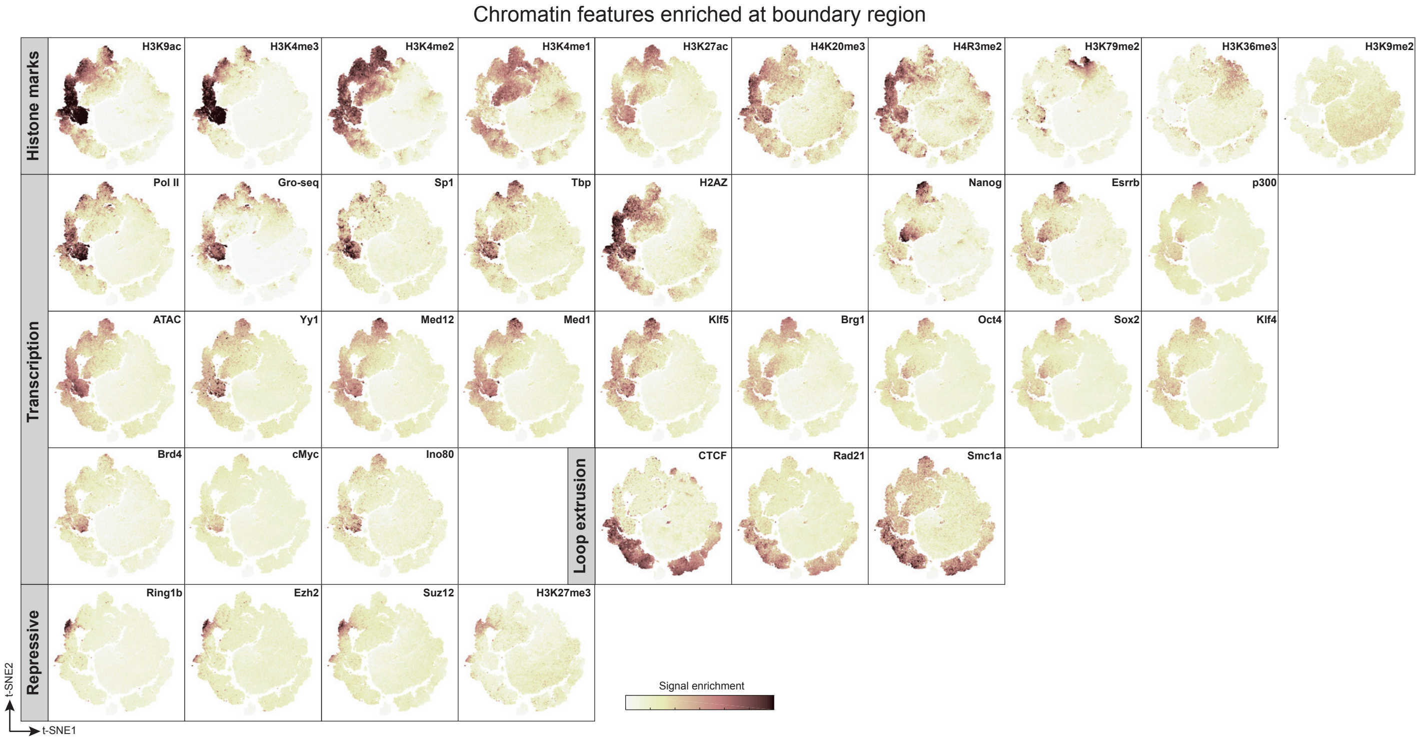

**fig. S6. PCA and t-SNE analysis of single boundaries.** (**A**) Principal Component Analysis of genome-wide data enriched at the microTAD boundaries. Y-axis shows the PCA coefficients (also known as loadings) for each component, and x-axis shows the observations. H3K4me3 and H3K9ac are the heaviest loadings for the first two components. Factors bound on cis-regulatory elements contribute to the second components. CTCF and cohesin are the most significant factors in PC3. PC1 – PC3 can explain ~75% of the variation. (**B**) A complete example of a t-SNE analysis of single microTAD boundaries. MicroTAD boundaries were plotted by t-SNE embedded points, and color-coded by the enrichment of genomic data for each boundary. Dimension-reduced data with the first ten PCA components (>90% of variance) were used for t-SNE analysis. Histone modifications on the boundary-flanking nucleosomes and CTCF/cohesin are sufficient to classify the main populations of the microTAD boundaries.

Supplemental Figure 7

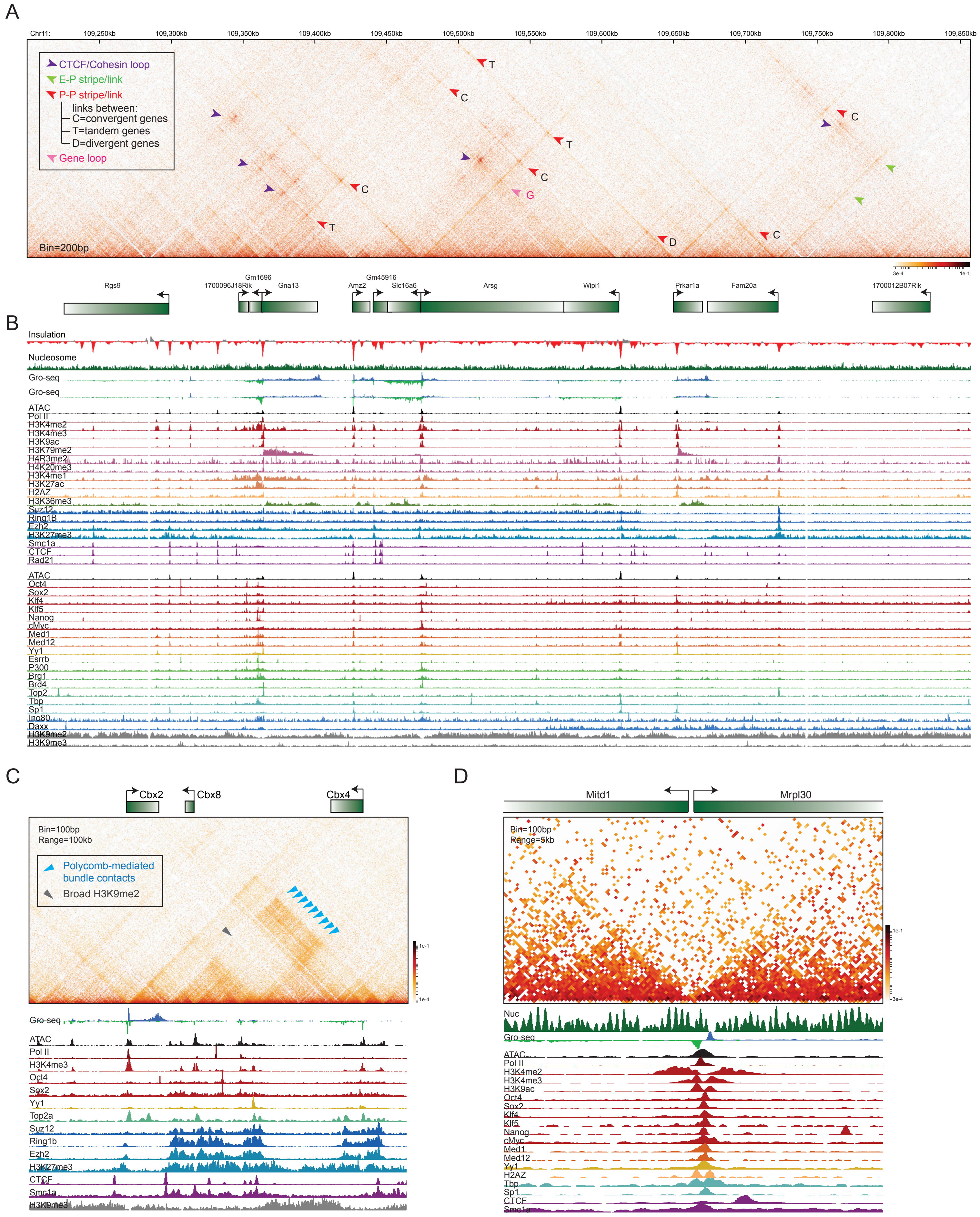

**fig. S7. Intricate chromatin organizations within TADs.** (**A-B**) Nucleosome-resolution chromatin organization in a 650-kb region in chr11. Contact map is overlaid with gene annotations and browser tracks of the relevant genomic data. Arrows highlight CTCF/cohesin-mediated loops (purple), E-P stripe/links (green), P-P stripe/links (red), and gene loops (pink). Orientations of P-P stripe/link are labeled as "C" for convergent genes, "T" for tandem genes, and "D" for divergent genes. The contact map demonstrates the nested chromatin architecture within a TAD. (**C**) An additional example of H3K27me3-associated chromatin bundles for Fig. 4C. Repressive chromatin interacts with another co-regulated repressive chromatin through an "interaction bundle" structure (blue arrows), rather than forming a loop-like conformation. (**D**) Example of promoter structure between a pair of divergent genes. The divergent promoter typically forms a strong boundary that is flanked by active nucleosomes. In some cases, the longer length of divergent promoters can be linked by P-P stripes (see example in (**A**), intergenic region between *Wipi1* and *Prkar1a* genes). We speculate this type of divergent P-P link might be mediated by antisense transcription.

### Supplemental Figure 8

A

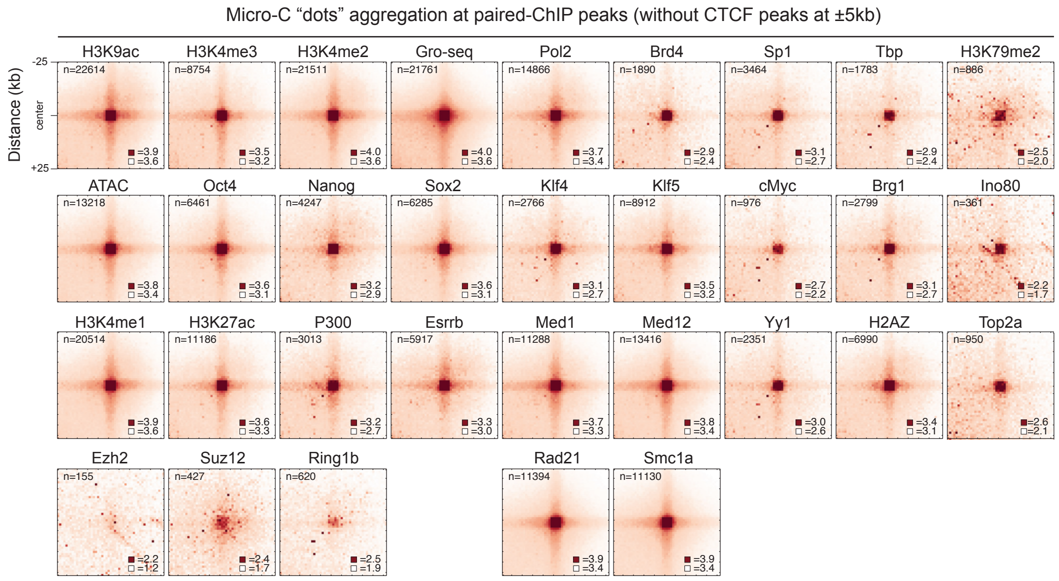

B

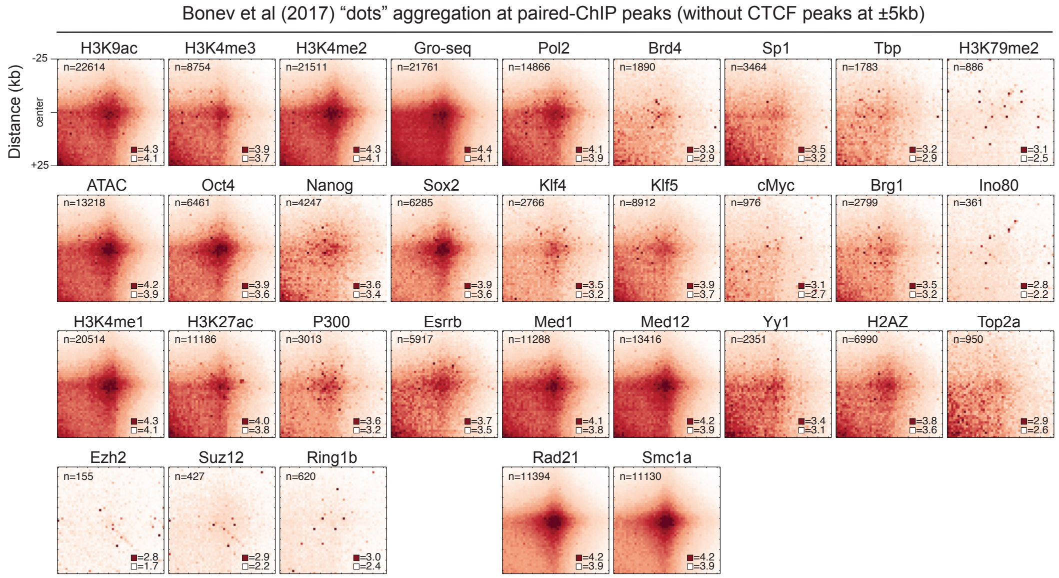

C

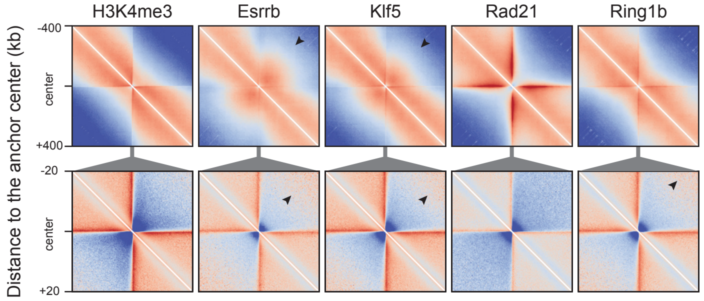

D

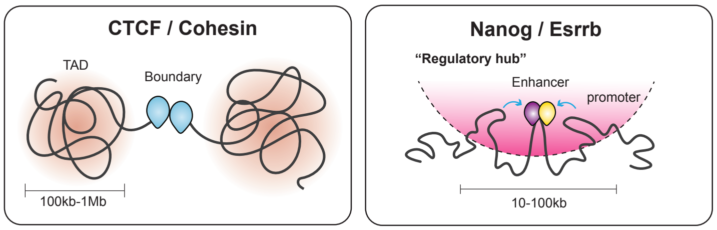

**fig. S8. Protein-centered chromatin links and hubs.** Pile-up analysis of dot enrichment was plotted according to the loci of paired ChIP-seq peaks. Only the pairs of ChIP-seq peaks shorter than 1-Mb distance were kept for the analysis. Loci also containing CTCF peaks within ± 5kb were removed. (**A**) Pile-up dot analysis with Micro-C data compared to (**B**) results by Bonev et al. (2017) Hi-C data (19). Micro-C presents a cleaner and sharper dot enrichment in all cases. Dots with weaker enrichments in the Hi-C data appear to relate to enhancer and repressive functions. (**C**) Additional examples of pile-up analysis of the target-centered chromatin structure for Fig. 4F-G. The top panel is the large scale of chromatin structure in a ±400-kb window and the bottom panel is a ±20-kb region zoomed in from the center of the top panel. Transcription associates with the strongest local boundary and stripes, and CTCF/cohesin contribute to the distal chromatin structures. (**D**) Schematics illustrate the proposed model of protein-centered chromatin structures. CTCF/cohesin and transcription factors form strong boundaries that spatially separate TADs and microTADs. Enhancer factors are concentrated in a local regulatory hub, which can mediate efficient enhancer-promoter contacts.

Supplemental Figure 9

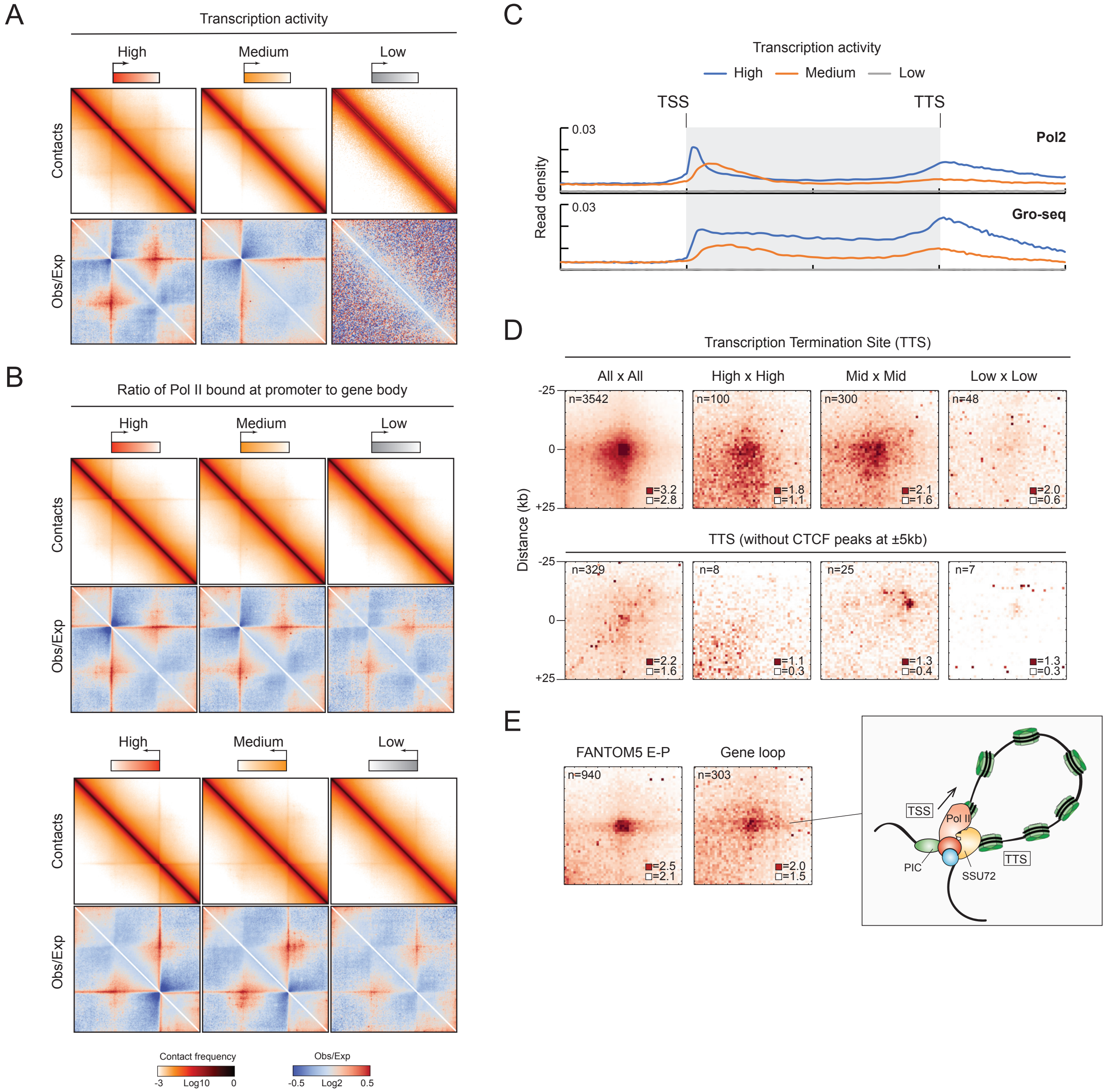

**fig. S9. Gene-centered chromatin structures.** (**A-B**) Rescaled pile-up analysis of gene folding. The contact matrix of each gene was rescaled to the same pseudo length and then aggregated according to the level sorted by transcription activity (A) or Pol II pausing ratio (B). Contact maps are shown in log10 contact counts normalized by matrix balancing or by distance. Consistent with the unscaled pile-up analysis in Fig. 5B, TSSs with high transcription rate show stronger local boundary strength, and more intra-gene interactions and gene stripes than weakly transcribed genes. Top and bottom panels in (B) highlight that gene orientation determines the orientation of the gene stripe. (**C**) Metagene analysis of Pol II and GRO-seq signal enrichment. Gene lengths were rescaled to the same size as shown in x-axis. Signal enrichments of each genome-wide data are plotted according to the transcription activity. (**D-E**) Pile-up analysis of dot enrichment between TTS, TSS-TTS within a gene (gene loop), and E-P links identified by FANTOM project (http://fantom.gsc.riken.jp/5/) (*26*). There is no significant enrichment of TTS links after removing CTCF peaks bound within ±5 kb, arguing that TTS might not contact another TTS via TTS-TTS looping in the absence of CTCF. Furthermore, we found moderate dot enrichment in gene loops and FANTOM5 E-P links. Gene loops might not be ubiquitous cases in mammalian cells, as we discovered 303 significant gene loops in mESC (FDR < 0.1). Also, there are only 940 (out of 47,971) E-P interactions that were identified by the FANTOM5 project are significantly enriched in Micro-C data. Schematic describes the current model of the gene looping structure. TSS and TTS of a gene are in contact with each other, which might be mediated by Pol II complex and SSU72 (RNA Polymerase II CTD Phosphatase)(*27*).

### Supplemental Figure 10

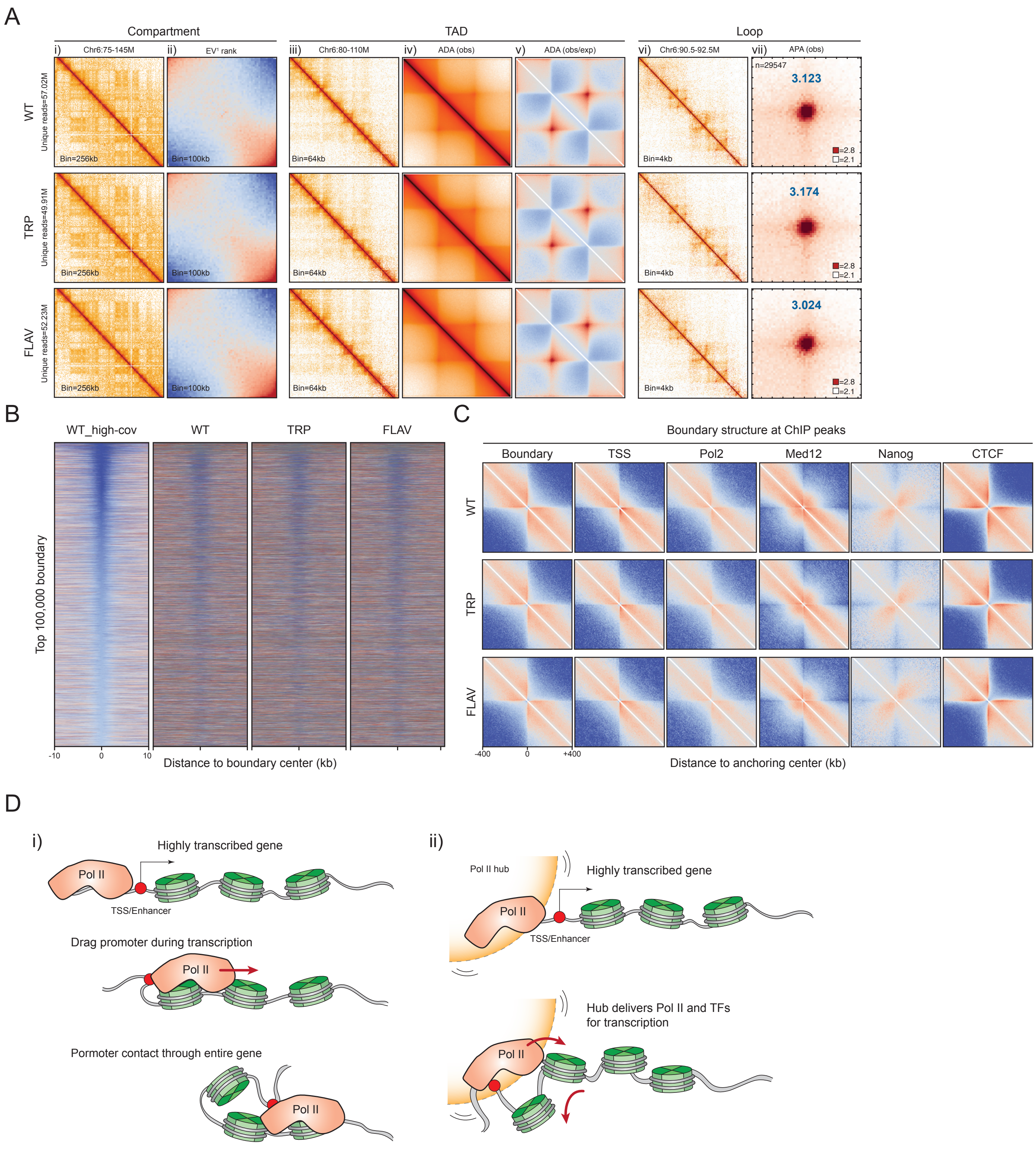

**fig. S10. Overview of the effect of Pol II inhibition on chromatin organization.** (**A**) Overview of chromatin organization upon Pol II inhibition. Compartments: i) An example of plaid-like chromosome compartments in Chr6 is shown as ICE balanced contact matrices at a 100-kb resolution. ii) Saddle plot for compartmentalization strength. Saddle plot was calculated by the average distance-normalized contact frequencies between 100-kb bins in cis with ascending eigenvector values ($EV^1$). The upper-left and bottom-right represent the contact frequency between B-B and A-A compartments and upper-right and bottom-left show the frequency of inter-compartment interactions. TAD: iii-iv) An example of TADs at 30M region in chr6. TADs were rescaled and aggregated at the center of the plot with ICE normalization or distance normalization. Loop: v) An example of loops in a 300-kb locus of chr6. vi) Pile-up analysis of loop enrichment. There is no significant change in these chromatin structures after Pol II inhibition for 45 minutes. Note that Micro-C with ~50M reads is capable of visualizing chromatin loops. (**B**) Pol II inhibition has little effect on boundary strength. Heatmaps were plotted by data with 1-kb resolution and centered at the top 10,000 microTAD boundaries flanked by ±10-kb region. (**C**) Pol II inhibition does not globally affect protein-centered chromatin structures. Pile-up analysis was centered at the query sites with ±400-kb window. (**D**) Schematics illustrate the proposed models of Pol II-mediated gene structures. In the model (i), Pol II and transcription machinery drag the promoter interacting through the entire gene body, resulting in the gene stripe structure in the Micro-C contact map. In the model (ii), the promoter is trapped within a hub that enriched with the Pol II complex, which can deliver essential components for gene transcription. This mechanism could also promote the formation of the gene stripe in the contact map.

### Supplemental Figure 11

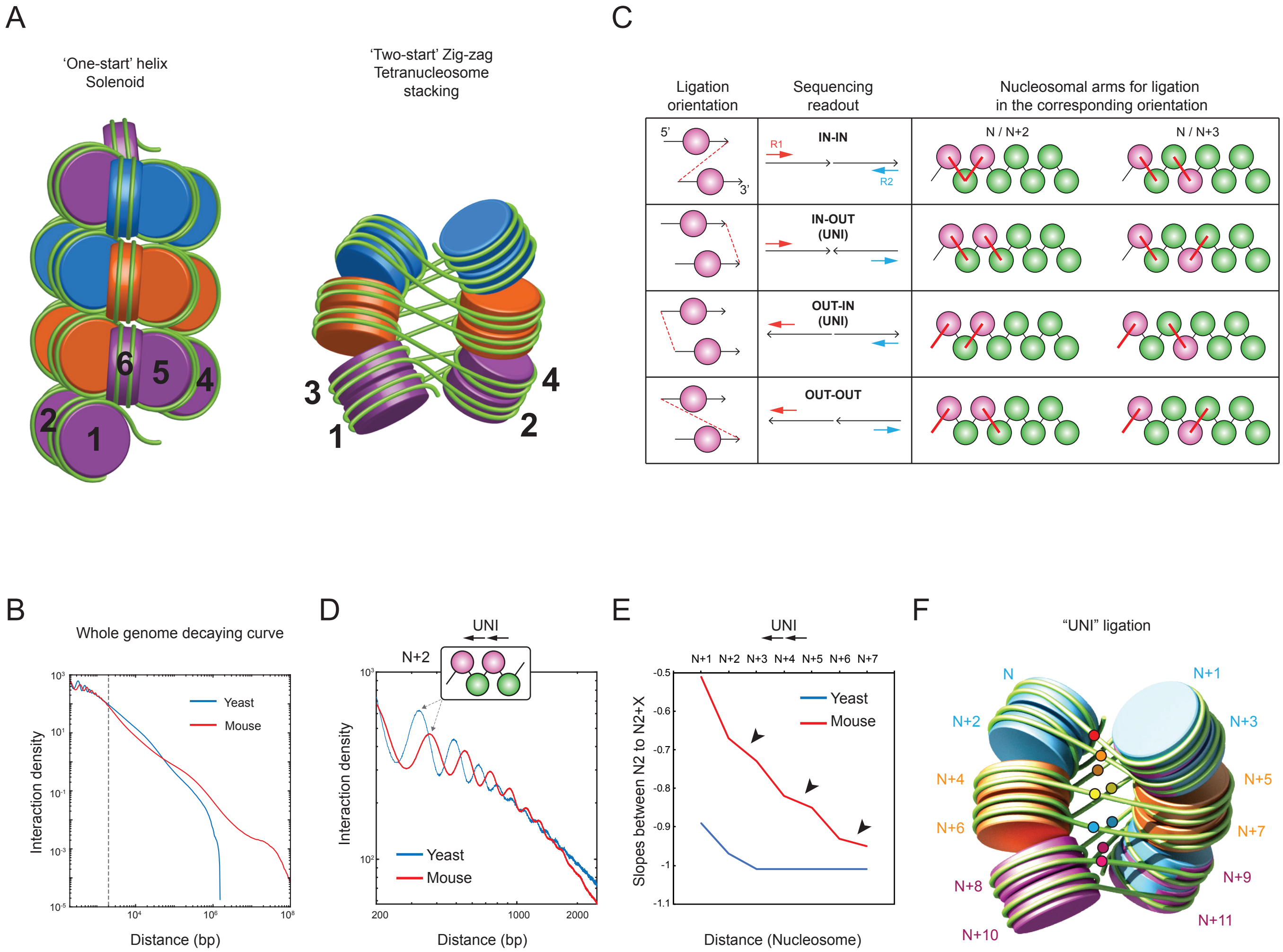

**fig. S11. 30 nm chromatin fibers.** (**A**) Schematics illustrate the models of 30 nm chromatin fiber. “One-start” solenoid model describes the nucleosome array follows a helical trajectory with ~6 nucleosomes per helical turn, which predicts that nucleosomes N and N+4/5 should to be in spatial proximity. “Two-start” zig-zag model describes the nucleosomes zigzag through a two-start helix (tetra-nucleosome unit) with a straight linker DNA so that alternate nucleosomes (N and N+2/3) are the interacting partners. The folding units are painted in the same color. (**B**) Decaying curves of mESCs and budding yeast BY4741 strain. X-axis is the distance in log10 bp, and y-axis is interaction density. The dashed line indicates the threshold of “rich” local nucleosome interactions, ~2000 bp. (**C**) Schematics describe the scenarios of nucleosome interactions. There are four orientations of nucleosome ligation. Dash line in red represents which nucleosome ends are ligated between adjacent nucleosomes. Based on the sequencing readout of the Micro-C products (labeled with R1 and R2), we separated the orientations to “IN-IN”, “IN-OUT”, “OUT-IN”, and “OUT-OUT”. Interactions in “IN-OUT” and “OUT-IN” are the same and decaying curves completely overlap. Thus, we combined them as “UNI” orientation. The last column on the table highlights which nucleosome ends are ligated in different orientations. (**D**) Interaction decaying curve of the “UNI” interactions at a distance from 200 to 2000 bp. Distances along the x-axis are from 200 to 2000 bp in log10 ratio, and y-axis shows the log10 contact density, normalized to ligated pairs per million reads per $bp^2$. (**E**) Slopes of N/N+X with distance. Curve indicates the slope between the peak point of N+2 and N+X in fig. S11D. X-axis represents the distances in units of nucleosomes, and y-axis is the slope of the indicated N/N+X. Black arrows highlight that every two adjacent nucleosomes (e.g., N+2 and N+3) have a similar level of interactions to nucleosome N. (**F**) The two-start zig-zag tetra-nucleosome stacks. The colored dots represent the ligated partners between N/N+X in the “UNI” orientation. Noted that “IN-IN” data was not included because undigested di-nucleosomes may bias our interpretation

Supplemental Figure 12

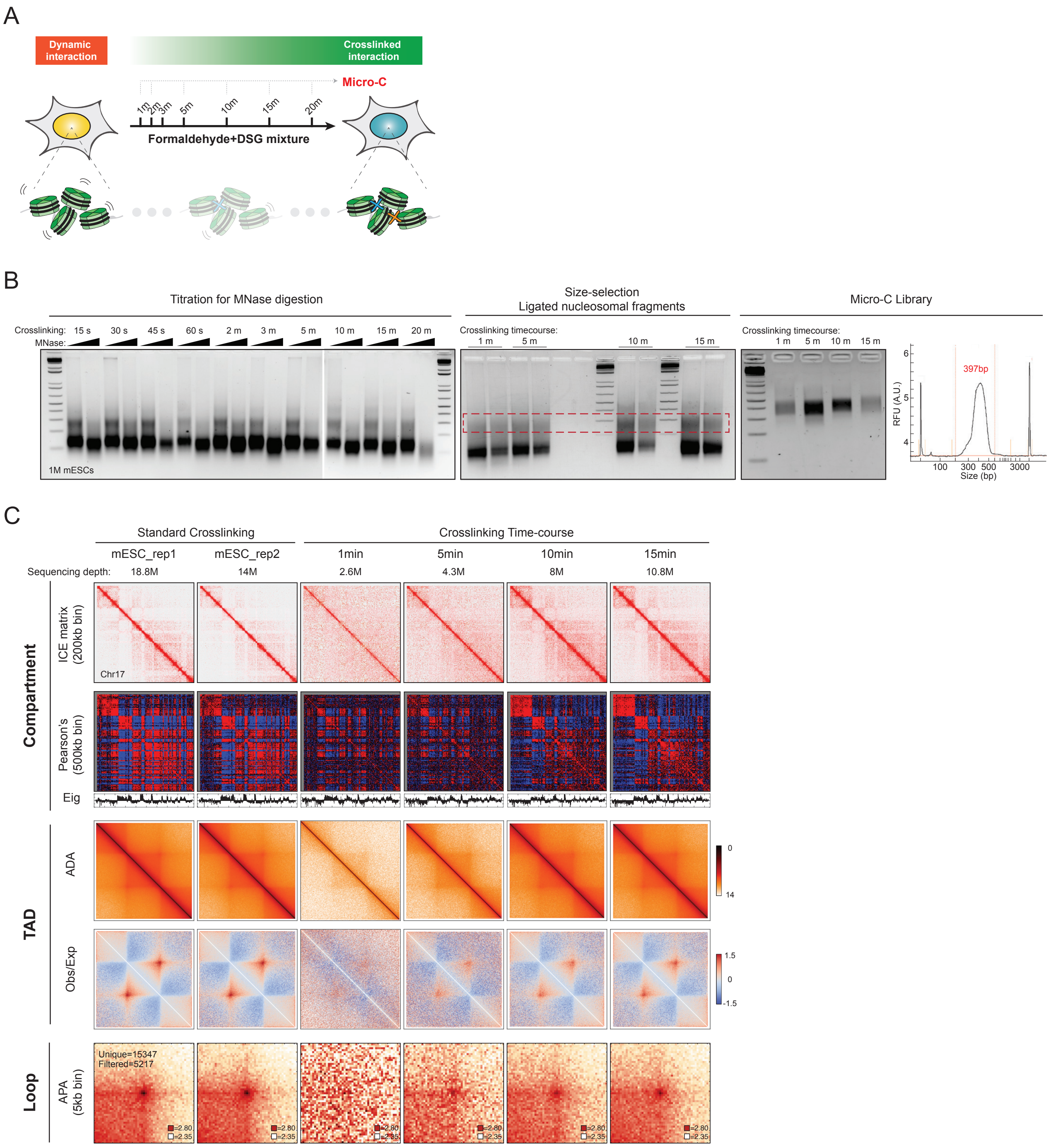

**fig. S12. Overview of crosslinking optimizations.** (**A**) Schematic describes the experimental design for crosslinking optimization. We collected fixed cells at a series of time for Micro-C experiments upon formaldehyde and DSG crosslinking. (**B**) First agarose gel image shows the titration of MNase digestion for each crosslinking time course. The time of crosslinking has little effect on MNase digestion. We then tested crosslinking efficiency by using the samples fixed for 1, 5, 10, and 15 minutes. The second gel image shows the size selection of the di-nucleosomal band. The third gel image and the bioanalyzer data indicate a sharp band at ~300-500 bp for the final sequencing library. (**C**) Overview of crosslinking-dependent Micro-C. Compartment: An example of plaid-like chromosome compartments in chr11 is shown as ICE balanced contact matrices, Pearson's correlation matrices, and Eigenvector analysis of the first principle component at 100-kb resolution. Compartment organizations become clear after 10 minutes of crosslinking. TADs: TADs were rescaled and aggregated at the center of the plot with ICE normalization or distance normalization. TAD structure can be clearly observed after 1 minute of crosslinking. Loop: Pile-up analysis of loop enrichment. Loops can be detected between 5 – 10 minutes of crosslinking. Note that the measurement cannot be directly interpreted as chromatin dynamics, given that crosslinking is a slow and inefficient process. We only use this data for the optimization of crosslinking conditions for the Micro-C protocol. Measuring real-time chromatin dynamics requires future investigations by single-molecule tracking studies.

#### Supplemental tables

| Samples | Description | GEO# | PMID# | Reference |
|---|---|---|---|---|
| ATAC-seq | Chromatin accessibility assay | GSE90892 | PMID: 28111071 | Chronis C, Fiziev P, Papp B, Butz S et al. Cooperative Binding of Transcription Factors Orchestrates Reprogramming. Cell 2017 Jan 26;168(3):442-459.e20. |
| Brd4 | Bromodomain containing protein | GSE87037 | PMID: 28472656 | Aldiri I, Xu B, Wang L, Chen X et al. The Dynamic Epigenetic Landscape of the Retina During Development, Reprogramming, and Tumorigenesis. Neuron 2017 May 3;94(3):550-568.e10. |
| Brg1 | Chromatin remodeler complex: SWI/SNF | GSE90893 | PMID: 28111071 | Chronis C, Fiziev P, Papp B, Butz S et al. Cooperative Binding of Transcription Factors Orchestrates Reprogramming. Cell 2017 Jan 26;168(3):442-459.e20. |
| Cbx3 | Heterochromatin protein | GSE44242 | PMID: 23748610 | Sridharan R, Gonzales-Cope M, Chronis C, Bonora G et al. Proteomic and genomic approaches reveal critical functions of H3K9 methylation and heterochromatin protein-1γ in reprogramming to pluripotency. Nat Cell Biol |
| Cbx5 | Heterochromatin protein | GSE64946 | PMID: 26415775 | Mattout A, Aaronson Y, Sailaja BS, Raghu Ram EV et al. Heterochromatin Protein 1β (HP1β) has distinct functions and distinct nuclear distribution in pluripotent versus differentiated cells. Genome Biol 2015 Sep 28;16:213. |
| cMyc | Pluripotency transcription factor | GSE90893 | PMID: 28111071 | Chronis C, Fiziev P, Papp B, Butz S et al. Cooperative Binding of Transcription Factors Orchestrates Reprogramming. Cell 2017 Jan 26;168(3):442-459.e20. |
| CTCF | Architectural protein / Loop extrusion factor | GSE90994 | PMID: 28467304 | Hansen AS, Pustova I, Cattoglio C, Tjian R et al. CTCF and cohesin regulate chromatin loop stability with distinct dynamics. Elife 2017 May 3;6. |
| Daxx | Histone chaperone for H3.3 | GSE70811 | PMID: 26340527 | He Q, Kim H, Huang R, Lu W et al. The Daxx/Atrx Complex Protects Tandem Repetitive Elements during DNA Hypomethylation by Promoting H3K9 Trimethylation. Cell Stem Cell 2015 Sep 3;17(3):273-86. |
| Esrrb | Enhancer factor | GSE90893 | PMID: 28111071 | Chronis C, Fiziev P, Papp B, Butz S et al. Cooperative Binding of Transcription Factors Orchestrates Reprogramming. Cell 2017 Jan 26;168(3):442-459.e20. |
| Ezh2 | Repressive chromatin factor | GSE85717 | PMID: 27783950 | Juan AH, Wang S, Ko KD, Zare H et al. Roles of H3K27me2 and H3K27me3 Examined during Fate Specification of Embryonic Stem Cells. Cell Rep 2016 Oct 25;17(5):1369-1382. |
| Gro-seq | Nascent RNA assay / Pol II pausing | GSE69143 | PMID: 26878240 | Flynn RA, Do BT, Rubin AJ, Calo E et al. 7SK-BAF axis controls pervasive transcription at enhancers. Nat Struct Mol Biol 2016 Mar;23(3):231-8. |
| H2AZ | H2A variant | GSE34483 | PMID: 23260488 | Hu G, Cui K, Northrup D, Liu C et al. H2A.Z facilitates access of active and repressive complexes to chromatin in embryonic stem cell self-renewal and differentiation. Cell Stem Cell 2013 Feb 7;12(2):180-92. |
| H3K27ac | Histone marks: enhancer | GSE90893 | PMID: 28111071 | Chronis C, Fiziev P, Papp B, Butz S et al. Cooperative Binding of Transcription Factors Orchestrates Reprogramming. Cell 2017 Jan 26;168(3):442-459.e20. |
| H3K27me3 | Histone marks: repressive chromatin | GSE90893 | PMID: 28111071 | Chronis C, Fiziev P, Papp B, Butz S et al. Cooperative Binding of Transcription Factors Orchestrates Reprogramming. Cell 2017 Jan 26;168(3):442-459.e20. |
| H3K36me3 | Histone marks: gene body | GSE90893 | PMID: 28111071 | Chronis C, Fiziev P, Papp B, Butz S et al. Cooperative Binding of Transcription Factors Orchestrates Reprogramming. Cell 2017 Jan 26;168(3):442-459.e20. |
| H3K4me1 | Histone marks: enhancer | GSE90893 | PMID: 28111071 | Chronis C, Fiziev P, Papp B, Butz S et al. Cooperative Binding of Transcription Factors Orchestrates Reprogramming. Cell 2017 Jan 26;168(3):442-459.e20. |
| H3K4me2 | Histone marks: transient? | GSE90893 | PMID: 28111071 | Chronis C, Fiziev P, Papp B, Butz S et al. Cooperative Binding of Transcription Factors Orchestrates Reprogramming. Cell 2017 Jan 26;168(3):442-459.e20. |
| H3K4me3 | Histone marks: active gene | GSE90893 | PMID: 28111071 | Chronis C, Fiziev P, Papp B, Butz S et al. Cooperative Binding of Transcription Factors Orchestrates Reprogramming. Cell 2017 Jan 26;168(3):442-459.e20. |
| H3K79me2 | Histone marks: DNA repair | GSE90893 | PMID: 28111071 | Chronis C, Fiziev P, Papp B, Butz S et al. Cooperative Binding of Transcription Factors Orchestrates Reprogramming. Cell 2017 Jan 26;168(3):442-459.e20. |

| | | | | |
|---|---|---|---|---|
| H3K9ac | Histone marks: active gene | GSE90893 | PMID: 28111071 | Chronis C, Fiziev P, Papp B, Butz S et al. Cooperative Binding of Transcription Factors Orchestrates Reprogramming. Cell 2017 Jan 26;168(3):442-459.e20. |
| H3K9me2 | Histone marks: heterochromatin / repeat elements | GSE54412 | PMID: 25637356 | Liu N, Zhang Z, Wu H, Jiang Y et al. Recognition of H3K9 methylation by GLP is required for efficient establishment of H3K9 methylation, rapid target gene repression, and mouse viability. Genes Dev 2015 Feb 15;29(4):379-93. |
| H3K9me3 | Histone marks: heterochromatin | GSE90893 | PMID: 28111071 | Chronis C, Fiziev P, Papp B, Butz S et al. Cooperative Binding of Transcription Factors Orchestrates Reprogramming. Cell 2017 Jan 26;168(3):442-459.e20. |
| H4ac | Histone marks: general active | GSE76760 | PMID: 26847871 | Gonzales-Cope M, Sidoli S, Bhanu NV, Won KJ et al. Histone H4 acetylation and the epigenetic reader Brd4 are critical regulators of pluripotency in embryonic stem cells. BMC Genomics 2016 Feb 4;17:95. |
| H4K20me3 | Histone marks: active/repressive? | GSE26680 | - | - |
| H4R3me2 | Histone marks: repressive? | GSE37604 | PMID: 24097435 | Girardot M, Hirasawa R, Kacem S, Fritsch L et al. PRMT5-mediated histone H4 arginine-3 symmetrical dimethylation marks chromatin at G + C-rich regions of the mouse genome. Nucleic Acids Res 2014 Jan;42(1):235-48. |
| Hira | Histone chaperone for H3.3 | GSE42152 | PMID: 24074864 | Banaszynski LA, Wen D, Dewell S, Whitcomb SJ et al. Hira-dependent histone H3.3 deposition facilitates PRC2 recruitment at developmental loci in ES cells. Cell 2013 Sep 26;155(1):107-20. |
| Ino80 | Chromatin remodeler complex: INO80 | GSE49137 | PMID: 24792115 | Wang L, Du Y, Ward JM, Shimbo T et al. INO80 facilitates pluripotency gene activation in embryonic stem cell self-renewal, reprogramming, and blastocyst development. Cell Stem Cell 2014 May 1;14(5):575-91. |
| Klf4 | Pluripotency transcription factor | GSE90893 | PMID: 28111071 | Chronis C, Fiziev P, Papp B, Butz S et al. Cooperative Binding of Transcription Factors Orchestrates Reprogramming. Cell 2017 Jan 26;168(3):442-459.e20. |
| Klf5 | Pluripotency transcription factor | GSE49848 | PMID: 24770696 | Aksoy I, Giudice V, Delahaye E, Wianny F et al. Klf4 and Klf5 differentially inhibit mesoderm and endoderm differentiation in embryonic stem cells. Nat Commun 2014 Apr 28;5:3719. |
| Med12 | General transcription factor | GSE22562 | PMID: 20720539 | Kagey MH, Newman JJ, Bilodeau S, Zhan Y et al. Mediator and cohesin connect gene expression and chromatin architecture. Nature 2010 Sep 23;467(7314):430-5. |
| Med1 | General transcription factor | GSE22562 | PMID: 20720539 | Kagey MH, Newman JJ, Bilodeau S, Zhan Y et al. Mediator and cohesin connect gene expression and chromatin architecture. Nature 2010 Sep 23;467(7314):430-5. |
| Nanog | Pluripotency transcription factor | GSE90893 | PMID: 28111071 | Chronis C, Fiziev P, Papp B, Butz S et al. Cooperative Binding of Transcription Factors Orchestrates Reprogramming. Cell 2017 Jan 26;168(3):442-459.e20. |
| Oct4 | Pluripotency transcription factor | GSE90893 | PMID: 28111071 | Chronis C, Fiziev P, Papp B, Butz S et al. Cooperative Binding of Transcription Factors Orchestrates Reprogramming. Cell 2017 Jan 26;168(3):442-459.e20. |
| P300 | Enhancer factor | GSE90893 | PMID: 28111071 | Chronis C, Fiziev P, Papp B, Butz S et al. Cooperative Binding of Transcription Factors Orchestrates Reprogramming. Cell 2017 Jan 26;168(3):442-459.e20. |
| Pol2 | Transcription | GSE58019 | PMID: 24999238 | Riising EM, Comet I, Leblanc B, Wu X et al. Gene silencing triggers polycomb repressive complex 2 recruitment to CpG islands genome wide. Mol Cell 2014 Aug 7;55(3):347-60. |
| Rad21 | Architectural protein / Loop extrusion factor | GSE90994 | PMID: 28467304 | Hansen AS, Pustova I, Cattoglio C, Tjian R et al. CTCF and cohesin regulate chromatin loop stability with distinct dynamics. Elife 2017 May 3;6. |
| Ring1b | Repressive chromatin factor | GSE77093 | PMID: 27941795 | Stanton BZ, Hodges C, Calarco JP, Braun SM et al. Smarca4 ATPase mutations disrupt direct eviction of PRC1 from chromatin. Nat Genet 2017 Feb;49(2):282-288. |
| Smc1a | Architectural protein / Loop extrusion factor | GSE123636 | 10.1101/495432 | Hansen AS, Hsieh THS, Cattoglio C, Pustova I, Darzacq X, Tjian R et al. An RNA-binding region regulates CTCF clustering and chromatin looping. BioRxiv 2018 Dec. |

| | | | | |
|---|---|---|---|---|
| Sox2 | Pluripotency transcription factor | GSE90893 | PMID: 28111071 | Chronis C, Fiziev P, Papp B, Butz S et al. Cooperative Binding of Transcription Factors Orchestrates Reprogramming. Cell 2017 Jan 26;168(3):442-459.e20. |
| Sp1 | General transcription factor | GSE52496 | PMID: 24850855 | Gilmour J, Assi SA, Jaegle U, Kulu D et al. A crucial role for the ubiquitously expressed transcription factor Sp1 at early stages of hematopoietic specification. Development 2014 Jun;141(12):2391-401. |
| Suz12 | Repressive chromatin factor | GSE39513 | PMID: 23101626 | Jia J, Zheng X, Hu G, Cui K et al. Regulation of pluripotency and self- renewal of ESCs through epigenetic-threshold modulation and mRNA pruning. Cell 2012 Oct 26;151(3):576-89. |
| Tbp | General transcription factor | GSE70661 | PMID: 27026076 | Langer D, Martianov I, Alpern D, Rhinn M et al. Essential role of the TFIID subunit TAF4 in murine embryogenesis and embryonic stem cell differentiation. Nat Commun 2016 Mar 30;7:11063. |
| Top2a | Toposiomerase | GSE45625 | PMID: 23698369 | Dykhuizen EC, Hargreaves DC, Miller EL, Cui K et al. BAF complexes facilitate decatenation of DNA by topoisomerase IIα. Nature 2013 May 30;497(7451):624-7. |
| WCE | Control | GSE90893 | PMID: 28111071 | Chronis C, Fiziev P, Papp B, Butz S et al. Cooperative Binding of Transcription Factors Orchestrates Reprogramming. Cell 2017 Jan 26;168(3):442-459.e20. |
| YY1 | Transcription factors / Architectural protein | GSE99518 | PMID: 29224777 | Weintraub AS, Li CH, Zamudio AV, Sigova AA et al. YY1 Is a Structural Regulator of Enhancer-Promoter Loops. Cell 2017 Dec 14;171(7):1573-1588.e28. |
| Chemical Nucleosome mapping | Nucleosome occupancy assay by chemical cleavage | GSE82127 | PMID: 27889238 | Voong LN, Xi L, Sebeson AC, Xiong B et al. Insights into Nucleosome Organization in Mouse Embryonic Stem Cells through Chemical Mapping. Cell2016 Dec 1;167(6):1555-1570.e15. |
| MPE-seq | Nucleosome occupancy assay by chemical cleavage | GSE69098 | PMID: 26080409 | Ishii H, Kadonaga JT, Ren B. MPE-seq, a new method for the genome-wide analysis of chromatin structure. Proc Natl Acad Sci U S A 2015 Jul 7;112(27):E3457-65. |
| MNase-seq | Nucleosome occupancy assay by MNase digestion | GSE69098 | PMID: 26080409 | Ishii H, Kadonaga JT, Ren B. MPE-seq, a new method for the genome-wide analysis of chromatin structure. Proc Natl Acad Sci U S A 2015 Jul 7;112(27):E3457-65. |
